# Resource allocation in the nodules of the *Pisum sativum - Rhizobium* symbiosis

**DOI:** 10.1101/2025.08.21.671492

**Authors:** Thomas J. Underwood, Philip S. Poole

## Abstract

Legumes host nitrogen-fixing bacteria, called rhizobia, within specialised root structures called nodules, where carbon from the plant is exchanged for ammonia fixed from N_2_ by the bacteria. Legumes can host multiple bacterial strains at the same time that vary in their fixation effectiveness but legumes sanction nodules containing less effectively fixing strains by reducing the provision of nutrients. Understanding how sanctions are applied by plants and how bacteria may try to avoid them is important to understand the stability of legume-rhizobial symbioses. Using near isogenic *Rhizobium leguminosarum* strains on pea we demonstrate that sanctions are sensitive to the proportion of nodules occupied by a less effective strain and by using split roots show that sanctions are applied based on global comparisons of nodules across the plants root system. By using several rhizobia with different levels of fixation, but all derived from the same parent, we show that pea plants can differentiate between bacteria with relatively small variations in fixation effectiveness. We demonstrate that peas integrate global and local signals in order to determine whether individual nodules are sanctioned. At the same time these results show that poorly fixing strains can avoid sanctions if they dominate nodulation.

## Introduction

Legumes, such as *Pisum sativum* (pea), can establish a mutualistic symbiosis with nitrogen-fixing bacteria known as rhizobia (Poole et al., 2018). In nitrogen-limited environments, this association allows the host plant to access a biologically usable form of nitrogen. Central to this symbiosis is the formation of root nodules, specialised organs within which rhizobia differentiate into Y-shaped bacteroids capable of fixing atmospheric nitrogen into ammonia (Mergaert et al., 2006). In exchange for fixed nitrogen, the host supplies carbon, primarily in the form of C4-dicarboxylates. This reciprocal exchange of nutrients underpins the mutualistic nature of the interaction (Udvardi & Poole, 2013).

These nodules provide a highly advantageous niche for rhizobia, enabling them to multiply to high densities before being released back into the soil upon nodule senescence (Timmers et al., 2000). However, this ecological opportunity opens the door for ‘cheater’ strains: rhizobia that colonise nodules and benefit from carbon and amplification without investing in nitrogen fixation (Denison, 2000).

Previous studies (Underwood et al., 2024; Westhoek et al., 2017, 2021) have demonstrated that built into this symbiosis on the part of the host plant is a sanctioning mechanism capable of punishing any ‘cheating’ bacteria that occupy a nodule. These ‘cheaters’ could potentially gain a fitness advantage by occupying a nodule, benefitting from the supply of carbon and amplification of numbers while expending less energy on nitrogen-fixation, thereby gaining a fitness advantage over a non-cheating strain.

Westhoek et al. (2021) showed that peas apply sanctions to nodules containing a strain that is less effective at fixing N_2_ by reducing their carbon supply. Remarkably, these sanctions are not applied to nodules based on a set threshold value of fixation. Instead, peas conditionally sanction a strain based on the other strains present. In this way a mediocre fixing strain may be sanctioned when a highly effective fixing strain is present but remains unsanctioned when a less effective strain is present.

This dynamic form of sanctioning raises important questions about the evolutionary stability of the symbiosis and the potential for exploitation by less cooperative strains. It has been shown that rhizobia of varying fixation effectiveness are present naturally (Berrada et al., 2019; Woliy et al., 2019), therefore, a strategy must exist through which less effective strains prosper. In this study, we investigated three possible avenues through which strains with reduced nitrogen fixation effectiveness might evade host sanctions and gain a competitive advantage:

1. High Proportion of Cheaters: In natural settings, rhizobial populations are unlikely to infect hosts in a 1:1 ratio. Since peas lack a strong ‘partner choice’ mechanism (Westhoek et al., 2017), they may be colonised by highly uneven ratios of effective and ineffectively fixing strains. If most nodules are occupied by low-fixing strains, stringent sanctioning may threaten the plant’s nitrogen supply. If, instead, the plant relaxes sanctions this may result in the increased abundance of the cheating strain. We tested this by inoculating peas with different ratios of a highly effective and a less effective strains and measuring sanction responses.
2. Local Clusters of Cheaters: Conditional sanctioning implies some form of comparison between nodules. Whether this occurs locally (between spatially proximate nodules) or globally (across the entire root system) remains unknown. If sanctions are applied based on local comparisons, a less effective strain might avoid punishment by clustering with similar nodules. To assess this, we employed a split-root system to spatially separate strains and evaluate nodule-specific responses.
3. Smaller Differences in Fixation Effectiveness: To date, studies have compared strains with relatively large differences in fixation capacity (Westhoek et al., 2021). However, it remains unclear how finely tuned is the ability of plants to discriminate between strains. If the resolution is coarse, a strain with only slightly reduced fixation could avoid sanctions. To test this, we introduced a third strain, a *fixL* double mutant (Rutten et al., 2021) which fixes nitrogen at a level intermediate to the previously tested Fix^+^ and Fix^int^ strains from Westhoek et al., (2021). This allowed us to assess the sensitivity of the sanctioning system of plants to subtle differences in symbiont performance.

Understanding how less effective or ‘cheating’ strains might evade host control is crucial for explaining the evolutionary dynamics of the legume rhizobia symbiosis and for maximising the continued benefit of the symbiosis to modern agriculture. The scattered, paraphyletic distribution of nodulation within the nitrogen-fixing clade of Rosids (Doyle, 2016), including partial losses within the Fabaceae, raises the possibility that uncontrolled symbiont exploitation may contribute to the loss of nodulation in some lineages.

Our findings demonstrate that peas modulate the severity of sanctions based on the relative abundance of strains, employing a global rather than a local mechanism of comparison. Additionally, peas can detect finer gradations in fixation effectiveness than previously shown, although an even smaller difference may still provide a loophole for cheating strains. These results refine our understanding of host control mechanisms and their limitations, with implications for the stability and breakdown of mutualism in this important symbiosis.

## Methods

### Rhizobial Strains and Culture conditions

The strains used in this work are all derived from an effective nitrogen-fixing bacteria Rhizobium leguminosarum bv. (Rlv) 3841 that is a root symbiont of *Pisum sativum* L. cv. Avola (pea) (Johnston & Beringer, 1975). The strains used varied in their fixation ability or fluorescent tag but are otherwise isogenic (Table 1). Some of the strains were tagged with a fluorescent marker (mCherry, GFP) to distinguish between the nodules formed by each strain (Table 1). Strains were maintained on tryptone-yeast (TY) agar (Beringer, 1974) with the required concentrations of antibiotics (Table 1). For longer-term storage a solution of TY with 15-20% glycerol was inoculated and then stored at -80°C. For inoculation of peas rhizobia were grown on a TY agar slope. The slopes were then washed using UMS and the number of bacteria present in the medium was determined by measuring OD600 using a Genesys 250 UV-Visible spectrophotometer. This solution was then diluted into a solution of approximately 1×10^7^ ml^-1^.

**Table 1:** Rhizobial strains. All strains derived from Rlv3841 and provided with a strain code, resistance markers, short description and reference.

| Name | Strain | Antibiotic resistance | Description | Reference |
| --- | --- | --- | --- | --- |
| Fix<sup>+</sup> | Rlv3841 | strep | Rlv3841 | (Johnston & Beringer, 1975) |
| Fix<sup>+</sup> GFP | OPS1339 | Strep, GM | Rlv3841 Tn7-Gm- GFP | (Westhoek et al., 2021) |
| Fix<sup>int</sup> mCherry | OPS2269 | Step, Spec, GM, | Fixint Tn7-Gm-mCherry (Rlv3841 mutant, ΩSpc cassette in promoter region of *nifA*) | (Westhoek et al., 2021) |
| Fix<sup>L</sup> | LMB496 | Strep, spec, | hfixL<sub>9</sub> :: omega spec, hfixL<sub>c</sub>:Pk19 | (Rutten et al., 2021) |

### Plant Growth

Peas were surface sterilised (1 minute in 95% ethanol followed by 5 minutes in 20% NaClO), rinsed with sterile water and left to germinate for five days on 1% w/v agar plates at room temperature in the dark.

#### Single pot experiments

After five days seedlings were transplanted to a sterilized 500 ml Azlon beaker for competition assays and 1 L Azlon beaker for Acetylene Reduction Assays. Beakers contained a 1:1 Mixture of silver sand and fine vermiculite and sterilized nitrogen-free nutrient solution (75 ml for 500 ml beakers and 150 ml for 1 L beakers). For single strain experiments 1×10^7^ cells of the desired rhizobial strain were added. For co-inoculated experiments the desired ratio (1:1 – 9:1) of the two rhizobial strains was added (0.5×10^7^ cells of each) (See Westhoek et al., (2017)). Beakers were covered with clingfilm to prevent aerial contamination. This was slit after a few days to allow seedlings to grow through. Plants were grown in a growth chamber (21 °C, 16 h photoperiod) for 28 days and watered as necessary from 7 days onwards.

#### Split root experiments

After five days the primary root of the seedling was cut laterally using a sterile scalpel, removing the tip of the root below where lateral root hairs had begun to form. These cut seedlings were then moved onto fresh 1% w/v agar plates, the edges of the plate were sealed with micropore tape and then placed in a growth chamber (21 °C, 16 h photoperiod) for a further five days. Seedlings were then placed across two 500 ml Azlon beakers with approximately half of the newly formed lateral roots placed into each beaker. The beakers contained a 1:1 Mixture of silver sand and fine vermiculite and 75ml of sterilized nitrogen-free nutrient solution. To each pot a total of 0.5×10^7^ rhizobial cells was added of the desired strain/strains. Plants were grown in a growth chamber (21 °C, 16 h photoperiod) for 28 days. The exposed root split across the pots was lightly watered every day for the first 7 days and then watered as necessary. In split root experiments a 1:1 ratio of each strain was always applied to the plant as whole.

### Harvesting

Plants were harvested 28 days post inoculation (dpi). Plants were removed from the sand vermiculite mix and washed carefully. Nodules were then imaged using a LEICA M165 FC fluorescent stereo microscope and an iBright FL1500 imaging system. Nodule occupants were identified based on their fluorescence (Fix^+^ tagged with GFP & Fix^int^ tagged with mCherry) or the lack of fluorescence (Fix^L^ untagged). The nodules of each strain were counted either per plant for single beaker experiments or per beaker for split root experiments.

### Nodule measurement

The five largest nodules of each strain on each plant or side of split root were selected by eye and then picked and photographed. These nodules were then measured using the Fiji analysis software (v2.14.0/1.54f). A ruler was included in each photograph of the nodules so that the scale for each image could be set by drawing a line of 10 mm over the markings of the ruler using a straight-line selection. Each nodule was then drawn around using the polygon selection tool and the area of the region of interest was measured. The mean of the five nodule sizes was calculated and taken as the average nodule size.

### Acetylene Reduction Assays

For acetylene reduction of experiments the whole plant was placed directly into a Schott bottle. The acetylene reduction assay was then carried out using the method described in Westhoek et al. (2021). Fixation effectiveness was then calculated as a percentage of the average fixation rate of Fix^+^.

### Statistical analysis

#### Fix^+^ vs Fix^int^ single pot co-inoculation

To test the correlation between the proportion of a strain within the inoculum and the proportion of nodules containing a strain a linear regression analysis was employed. A linear regression was also used to assess the correlation between the number of nodules a strain occupied and the average size of the nodules containing that strain.

#### Fix^+^ vs Fix^int^ split root co-inoculation

When testing for a difference in average size between Fix^+^ & Fix^int^ nodules in a split root system a nested linear mixed effect model was used in which the experiment number was a random factor and the plant number was used as a grouping factor allowing a pairwise comparison of nodules on the same plant across our biological replicates. As in the single pot experiments, when assessing the correlation between the number of nodules a strain occupied and the average size of those nodules, a linear regression was used

To test for a significant difference in the size of nodules on a split root plant when inoculated into separate pots or mixed into both pots a nested linear mixed effects model was used with experiment number as a random factor and plant as a grouping factor for pairwise comparison. This model was then tested using a one-way ANOVA, if the ANOVA showed a significant difference between groups this was then followed up with a Tukey’s post-hoc test.

#### Fix^+^, Fix^int &^ Fix^L^

To test the effect of co-inoculum on three strains of varying fixation effectiveness a linear model was produced for each strain comprised of the size of the nodules containing the strain and the co-inoculum (either one of the other two strains or the same strain). A one-way ANOVA was then used to test whether co-inoculum led to a significant variation in nodule size. If this test returned a significant difference, then a Tukey’s post-hoc test was used to compare each of the co-inoculum to the others.

All statistical analysis was carried out using R version 4.4.1 (2024-06-14)) and RStudio (2022.02.3+492 “Prairie Trillium”). Graphs were produced using Graph Pad version 9. An R markdown document is provided (Fig. S1). The model assumptions of equal variance and normality were assessed by visual inspection of residual plots (see R Markdown). Values from statistical tests are given where a significant trend or difference was found (p < 0.05). Where no significant trend or difference was found (p > 0.05) the values are not given. All statistical test outputs are provided in the R Markdown (Fig. S1). Data for each experiment is provided in Table S2.

## Results

### The severity of sanctioning is dependent on the proportion of cheats

To maximise the acquisition of nitrogen from nodule fixation legumes punish cheating strains. However, when the ratio of cheating to effective strains is varied it may be hypothesised that in order to still meet the plants nitrogen requirements it would be advantageous for the legume to relax sanctions as the proportion of cheaters increases. By the same logic it would also be expected that sanctions would become more severe as the proportion of cheaters decrease.

To test the effect of varying the frequency of the cheating strain, peas were inoculated with Fix^+^ (GFP) and Fix^int^ (mCherry) at varying ratios (1:9, 1:2, 1:1, 2:1, 9:1). As shown in (Westhoek et al., 2017) there was a significant linear correlation between a strains proportion of the inoculum and the proportion of nodules occupied by the strain (y = 0.99X – 1, R^2^ = 0.961, p < 0.0001)(Fig. 1 A). The average size of the nodules containing the less effective Fix^int^ was significantly positively correlated with the frequency of nodules containing Fix^int^ (y = 30X – 16, R^2^ = 0.731, p < 0.0001)(Fig. 1 B). In contrast, the size of nodules containing the more effective Fix^+^ strain did not show a significant correlation to the number of nodules the strain occupied (Fig.1 C).

**Fig 1.**
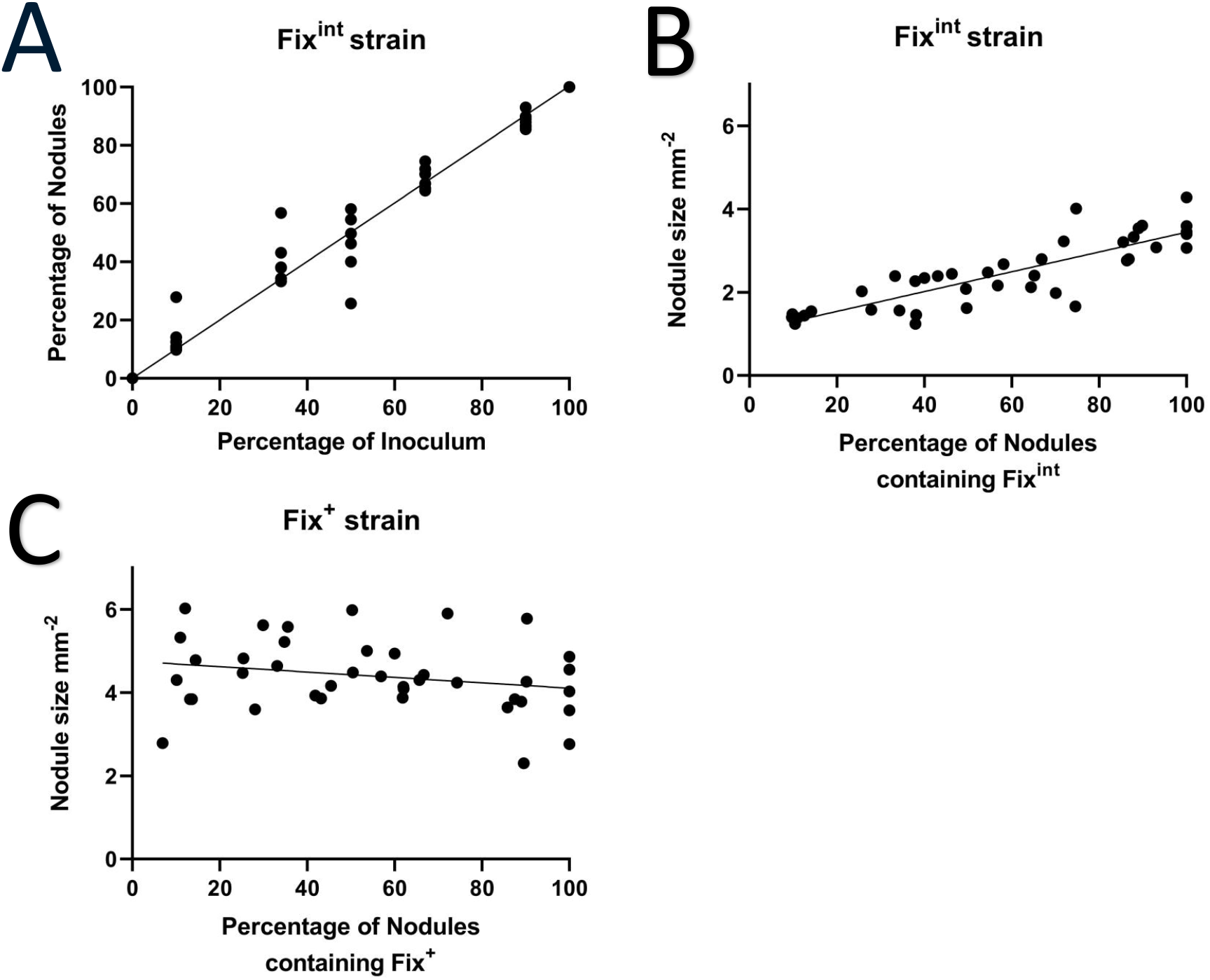
The sanctioning of a less effective nodule is dependent on their proportion of total nodules. For peas inoculated with a Fix^+^ and Fixint strain, the percentage of nodules containing a Fix^int^ strain and the percentage of the Fix^int^ strain within the inoculum applied (A). The average size of the Fix^int^ (B) and Fix^+^ (C) containing nodules on these plants plotted against the percentage of nodules containing the relevant strain. Each point represents the percentage (A) or the average size (B & C) for one plant. Trend line is a linear regression.

From this result it may be concluded that, as hypothesised, the severity of sanctions imposed on a less effective strain varies with the strains relative frequency. At the same time this was not true for the more effective strain. This suggests that the more effective strain is always receiving the maximum benefit from the host plant and the legume will then vary its treatment of the less effective strain to maximise its supply of nitrogen from nodules.

### Sanctioning is based on a global comparison between nodules

To test to what extent sanctioning is controlled through a global or local signalling mechanism a split root system for pea was developed, allowing the root system of a single pea plant to be inoculated with two different strains in physically separate pots thereby eliminating any potential effect of a local signalling mechanism between nodules that are proximal to each other.

When spatially separated Fix^+^ nodules were significantly larger than the Fix^int^ nodules (est. = 3.4191, SE = 0.1717, df = 9, t = 19.909, p < 0.0001) (Fig. 2A). When comparing the size of the Fix^int^ nodules with the proportion of nodules occupied by Fix^int^ these plants also showed the same pattern seen in the single pot experiments i.e. as the proportion of the less effective strain increased so did the size of the nodules containing the less effective strain (y = 13X + 26, SE = 3.912, R^2^ = 0.531, p = 0.0101) (Figure 2B).

**Fig 2.**
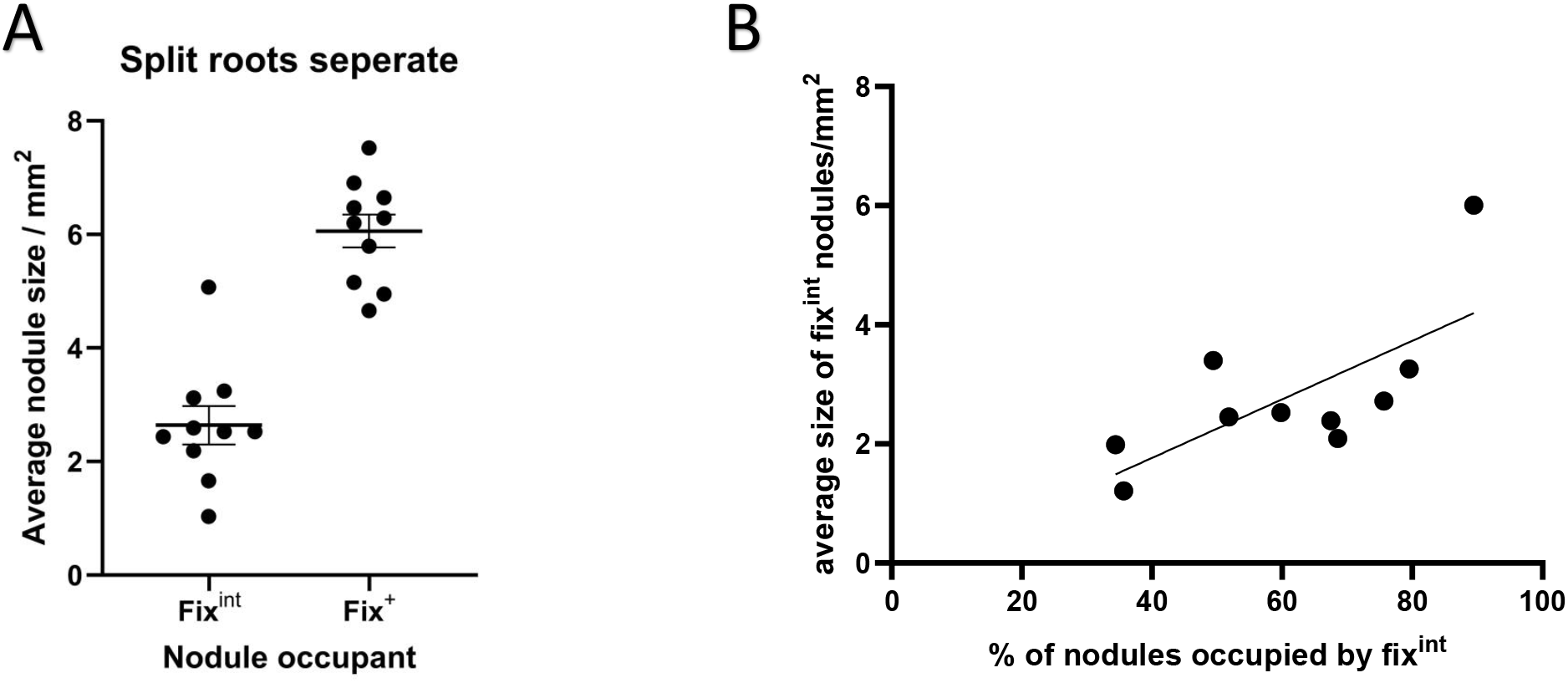
Split root peas still impose sanctions. Peas with split roots placed across two pots were inoculated with a Fix^+^ strain and a Fix^int^ strain with each strain added to a separate pot. The average size per plant of the nodules for each strain was measured (A). The relationship between the proportion of nodules occupied by the less effective Fix^int^ and the size of those nodules was also compared (B). Each point indicates the average value for a single plant. For B the trend line is a linear regression.

This result demonstrates that the effect of sanctions is not altered by spatial separation of the nodules. Therefore, legumes carry out a global comparison between the nodules present and apply sanctions accordingly

### Proximity does not alter sanctioning

It is conceivable that there is a local as well as a global element to the application of sanctions. To test this a series of split root plants were inoculated with both the Fix^+^ and Fix^int^ strain into both pots. These mixed rather than separately inoculated split root plants could then be compared to the separately inoculated plants. If there was a proximity effect on sanctioning it would be expected that the severity of sanctioning would significantly vary between these two sets of split root plants.

The Fix^int^ nodules on split roots with mixed inoculum were significantly smaller than the Fix^+^ nodules on the same plants (estimate = 3.2811, SE = 0.1642, df = 8.99, t = 19.98, p < 0.0001) (Figure 3A). When comparing the nodules on the separately inoculated split roots to those on the mix inoculated plants there was no significant difference between the two sets of Fix^+^ nodules or the two sets of Fix^int^ nodules. As was the case for both the single pot and the separately inoculated split root plants the size of the Fix^int^ nodules was significantly correlated with the proportion of nodules occupied by the Fix^int^ strain (y = 14.2X + 25, SE = 4.783, R^2^ = 0.465, p = 0.018) (Figure 3B). In short, the results of the mix inoculated split root plants were identical to those of the separately inoculated split roots.

**Fig 4.**
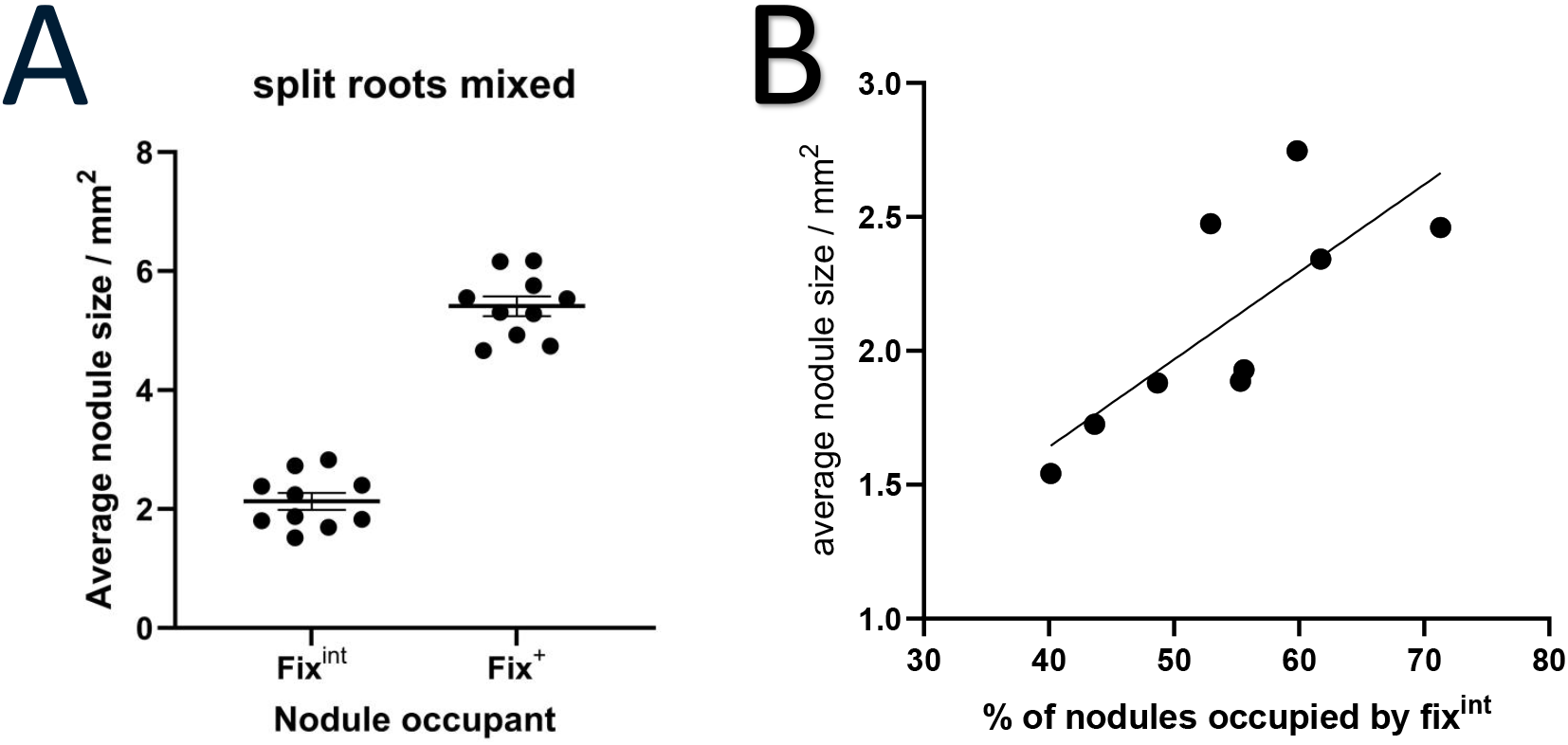
Sanctions are not modified by proximity. Peas with split roots placed across two pots were inoculated with a 1:1 mix of a Fix^+^ strain and a Fix^int^ strain added to each pot. The average size per plant of the nodules for each strain was measured (A). The relationship between the proportion of nodules occupied by the less effective Fix^int^ and the size of those nodules was also compared (B). Each point indicates the average value for a single plant. For B the trend line is a linear regression

These results demonstrate that the treatment of neither the more nor the less effective strain on a split root plant varies significantly due to the proximity of nodules containing a different strain. We therefore conclude that sanctioning is regulated by a global signalling mechanism that is unaffected by local cues.

### Peas can detect smaller changes in fixation effectiveness

To test the ability of peas to detect smaller differences in fixation effectiveness plants were inoculated with one of three combinations of strains. The Fix^+^ & Fix^int^ strains used in Westhoek et al., (2021) and Underwood et al., (2025) as well as each of these strains with the Fix^L^ strain from Rutten et al., (2021). This Fix^L^ strain fixes at a rate in between that of the Fix^int^ and Fix^+^ strain (Fig. S3) and therefore the differences in fixation between the Fix^L^ strain and the others is smaller than between Fix^+^ and Fix^int^.

When Fix^L^ was co-inoculated with Fix^+^ the nodules containing Fix^L^ were significantly smaller than when Fix^L^ was the sole inoculant (estimate = -1.889, 95% CI = -3.518: -0.258, p = 0.023) or when Fix^L^ was co-inoculated with Fix^int^ (estimate = -2.450, 95% CI =) (Fig. 4A). Nodules containing the Fix^int^ strain were significantly smaller when co-inoculated with Fix^+^ (estimate= = -2.136, 95% CI = -4.010: -0.889, p = 0.003) or Fix^L^ (estimate= = -1.516 95% CI = -2.807: -0.226, p = 0.021) compared to when Fix^int^ was the sole inoculant (Fig. 4B). Fix^+^ nodules did not significantly vary in size regardless of co-inoculant or when solo inoculated (Fig. 4C).

**Fig. 4.**
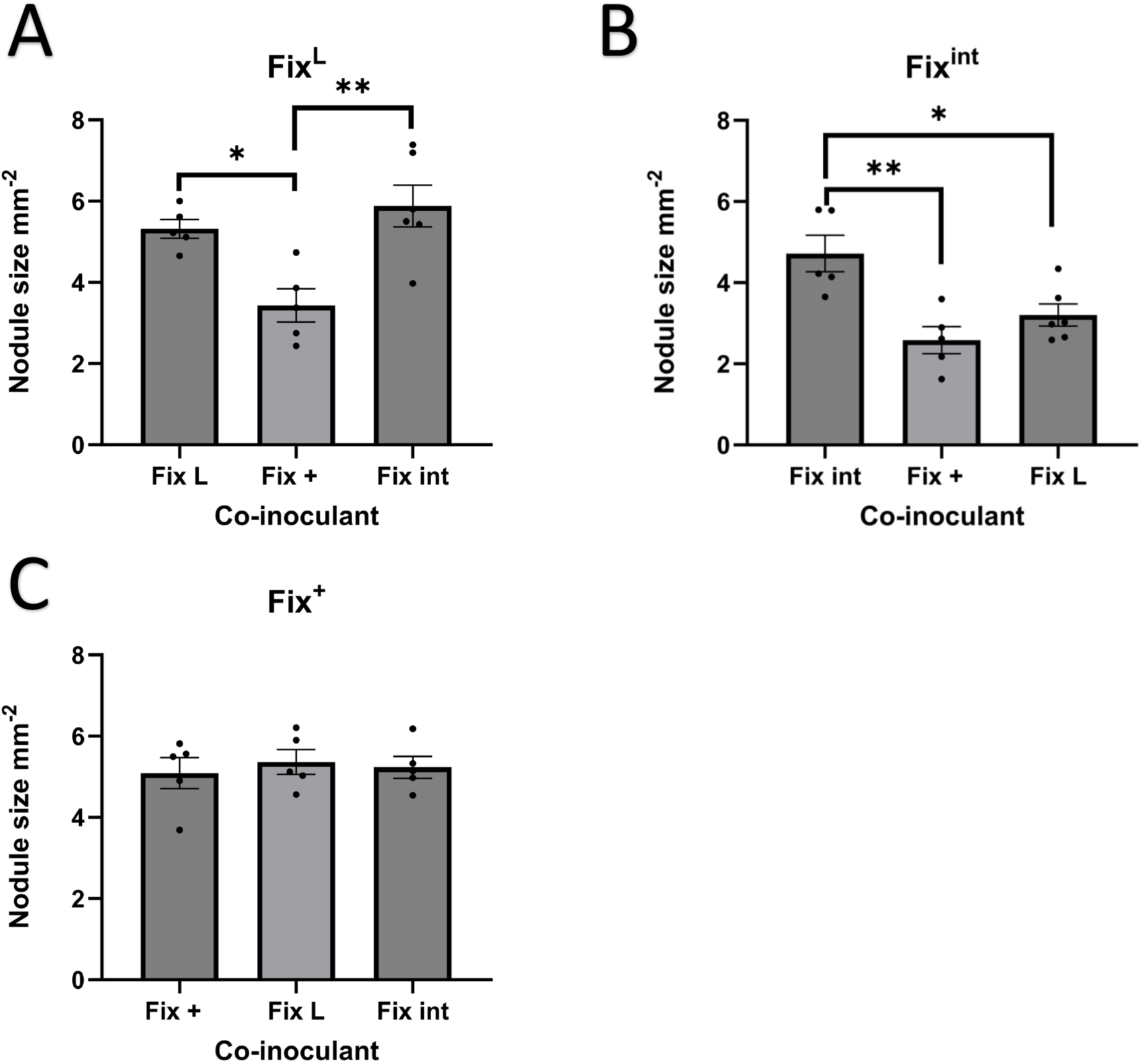
Peas can detect small variations in fixation and apply sanctions accordingly. Peas were inoculated with a Fix^L^ (A), Fix^int^ (B) or a Fix^+^ (C) strain. These strains were co-inoculated either with themselves or with one of the other two strains. Nodules were harvested after 28 days and average nodule size was measured. Each black point indicates the average nodule size for one plant. Bars indicate the mean of the average nodule sizes. Error bars indicate the standard error of the mean. Significant pairwise comparisons from Tukey’s post hoc test are given. * = p< 0.05, ** = p< 0.01.

The sanctioning of Fix^int^ when co-inoculated with Fix^L^ and the sanctioning of Fix^L^ when co-inoculated with Fix^+^ indicates that the host plant was capable of differentiating between the three strains and successfully punishing the less effective strain.

## Discussion

We have shown that peas moderate the severity of the sanctions imposed on a less effective strain depending on the proportion of nodules occupied (Fig. 1). We also conclude that peas compare nodules across their root system through a global signalling mechanism (Fig. 2) that does not include a local comparison between nodules that are proximal to each other (Fig. 3). We have tested the ability of peas to differentiate between smaller variations in fixation effectiveness than has been previously tested and found that they were still able to sanction the less effective strain (Fig. 5). However, this result does not allow us to make a broader conclusion on the ability of a less effective strain with a smaller variation in fixation effectiveness to evade sanctions.

This study has tested three avenues for cheating strains to prosper in a diverse population of nitrogen-fixing bacteria and has concluded that a viable avenue does exist. If a cheating strain dominates nodulation, then the plant will reward the strain. This is likely because when such a large number of nodules are occupied by a less effective strain the plant is unable to meet its nitrogen requirements by only rewarding the more effective strain. Thereby forcing the plant to reduce or entirely suspend sanctioning of the less effectively fixing strain. This result partially explains the inability of conditional sanctioning to eradicate less effective strains in wild populations. When considering the use of effectively fixing inoculum to boost yields in an agricultural setting it is therefore crucial to consider the relative size of the population and the competitive nodulating ability of the strain applied relative to the pre-existing strains in the soil. If too low a population of the introduced strain is present or if the strain cannot nodulate competitively then too few nodules will be occupied by the effective fixing strain allowing less effectively fixing strains to evade sanctions removing the potential yield benefits from the applied strain.

From the results of this study we can draw conclusions about the mechanism that underlies sanctioning. It is now clear that the first step in the application of sanctions is for the plant to comprehensively detect and then integrate information about the global nitrogen status. This involves detecting the individual output of all existing nodules, this must be the case as we have shown that plants carry out sanctioning based on a global comparison. Given the variation in sanction severity based on the proportion of nodules occupied by a strain it must also be concluded that the plant mediates sanctions based on the number of nodules present, their fixation rates and the plants nitrogen requirements. In short, the plant assesses how many nodules it needs to invest in to meet its nitrogen requirements. Not only this but our results show that the comparison between nodule fixation rates occurs at an even finer resolution than previously shown. This remarkable detection and information integration then results in individual nodule sanctioning outcomes.

A potential mechanism therefore requires a global signalling system that outputs developmental changes in the root architecture based on nitrogen detection. Candidate signalling molecules therefore include the plant hormones, auxin, and abscisic acid (ABA). Auxin has been shown to play a crucial role in nitrate signalling (Vega et al., 2019) where nitrate modulates auxin transport and accumulation. This process in part occurs via the action of the nitrate transceptor NRT1.1 (Krouk et al., 2010) which has already been hypothesised to play a regulatory role in sanctioning (Underwood et al., 2024). The hormone ABA is also a promising candidate for a signalling molecule in the application of sanctioning as it has been shown that nitrate stimulates ABA accumulation (Ondzighi-Assoume et al., 2016) while at the same time ABA plays a key role in in root architecture development (Signora et al., 2001). The role of ABA may be as a negative regulator of nodule sanctions as the presence of ABA inhibits expression of ABI1 which is a negative regulator of Snrk1 (Rodrigues et al., 2013) which is required for nodule malate production through the positive regulation of the genes MDH1 & 2 (Guo et al., 2023). This theory is further supported by the accumulation of ABA when nitrate is present (Ondzighi-Assoume et al., 2016) therefore it may be expected that ABA will accumulate around effectively fixing nodules.

This study demonstrates the extraordinary complexity of the regulatory mechanism of sanctioning as the plant must assess a myriad of factors to make individualised decisions for each nodule to maximise host benefit. Future studies must now look to elucidate the details of the genetic hormonal pathways that allow this incredibly intricate decision making to occur.

## Acknowledgements

This work was supported by the Biotechnology and Biological Sciences Research Council [BB/T008784/1].

