## Supplement figure 3 for "Resource allocation in the nodules of the *Pisum sativum - Rhizobium* symbiosis"

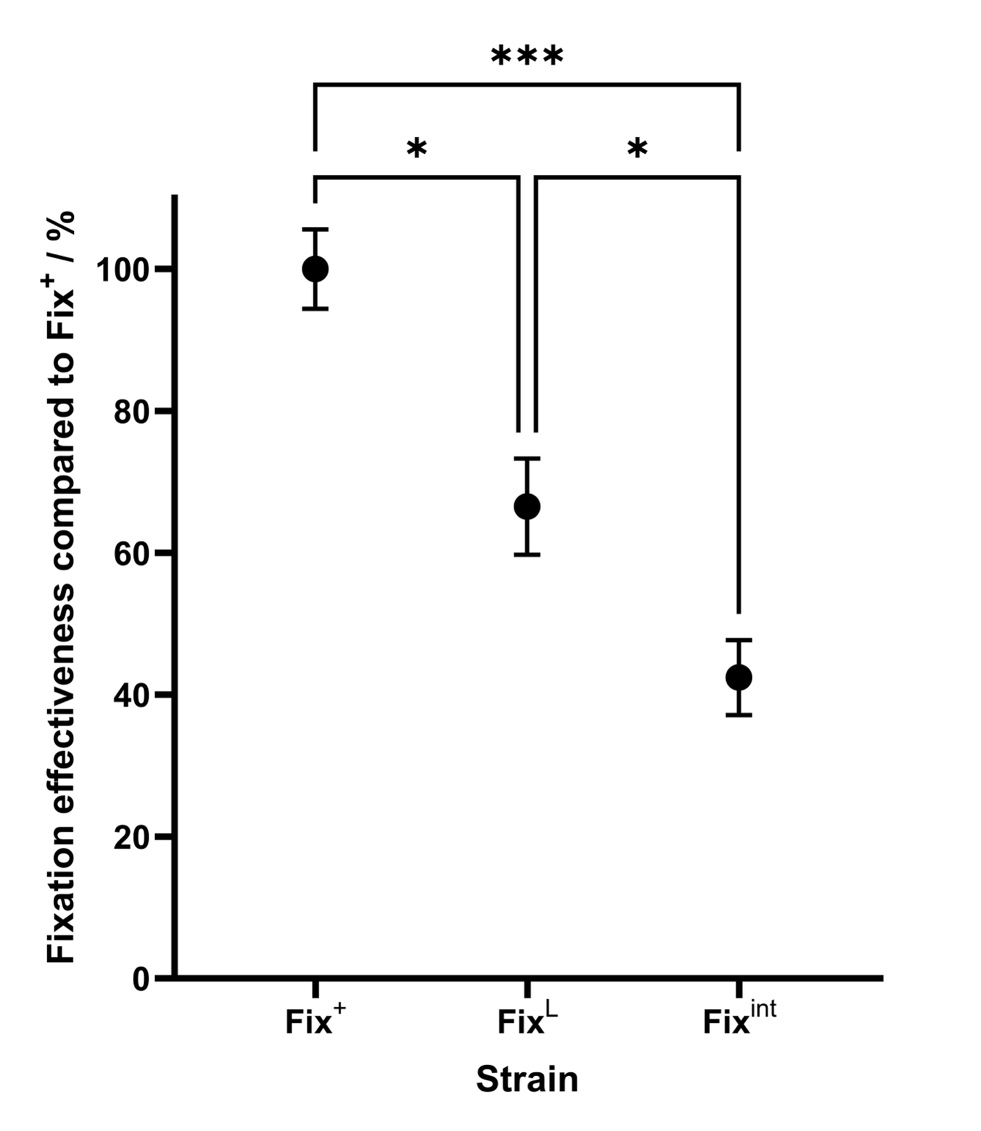

**Figure S1: Fixation effectiveness of three near isogneic strains**

The rate of fixation was measured for three strains. The values were converted to a percentage of the average fixation rate of Fix^+^ (wild-type Rlv 3841). Black dots denote average fixation effectiveness. Error bars are one standard error of the mean. Statistical comparison carried out though one-way ANOVA and Tukey’s post-hoc test. * = p < 0.05, *** = p < 0.001.
