## Supplementary figure 1 for "Resource allocation in the nodules of the *Pisum sativum - Rhizobium* symbiosis"

### ISME

2025-08-20

The first question we sort to answer was what effect would changes to the proportion of cheating vs non cheating strains have on host sanctions

```
#We compared the proportion of nodules occupied by fix int with
#the numbers of nodules occupied by fix int

proportion_model_inoculum <-
  lm(Proportion_data$`Inoculum int` ~ Proportion_data$`percent int`)
#This model is for Fix int nodules compared with the inoculum

#We then compared the size of nodules to
#the proportion of nodules occupied by that strain

proportion_model_size_int <-
  lm(Proportion_data$`percent int` ~ Proportion_data$`int av`)
#this model is for int nodules

proportion_model_size_plus <-
  lm(Proportion_data$`percent plus`~Proportion_data$`Plus av`)
#this is for plus nodules

plot(proportion_model_inoculum)
```

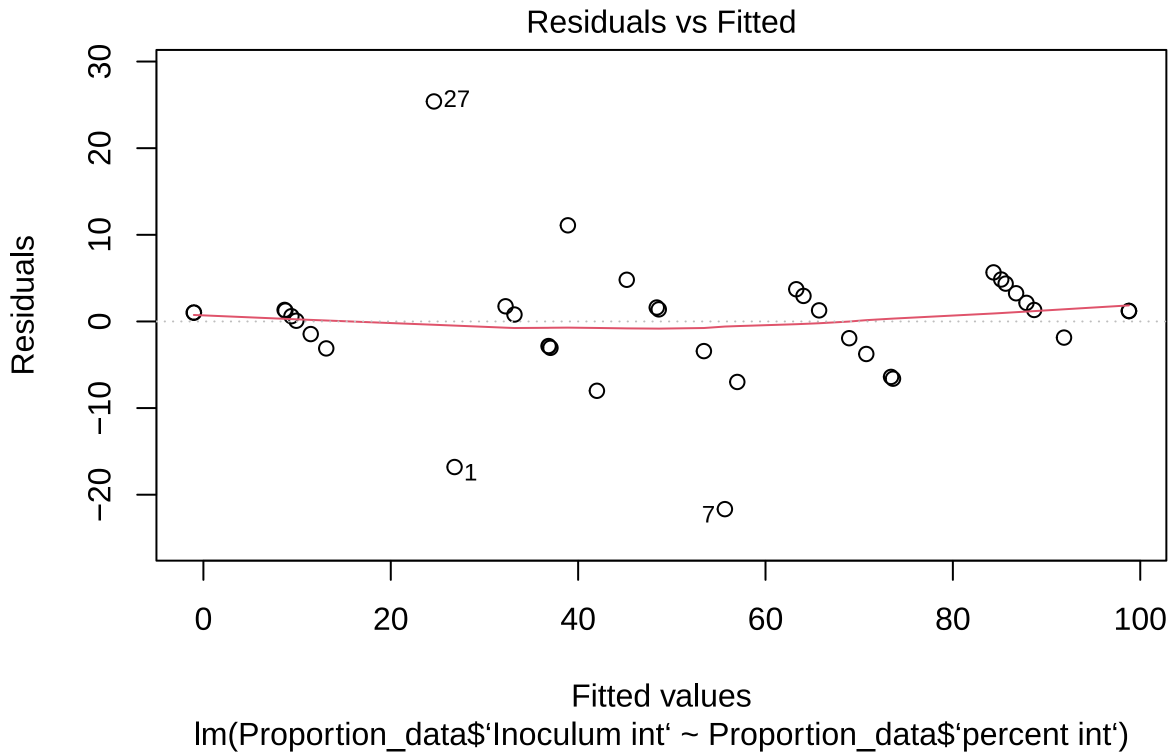
Residuals vs Fitted
Residuals
30
20
10
0
−10
−20
27
1
7
0
20
40
60
80
100
Fitted values
lm(Proportion_data$‘Inoculum int‘ ~ Proportion_data$‘percent int‘)

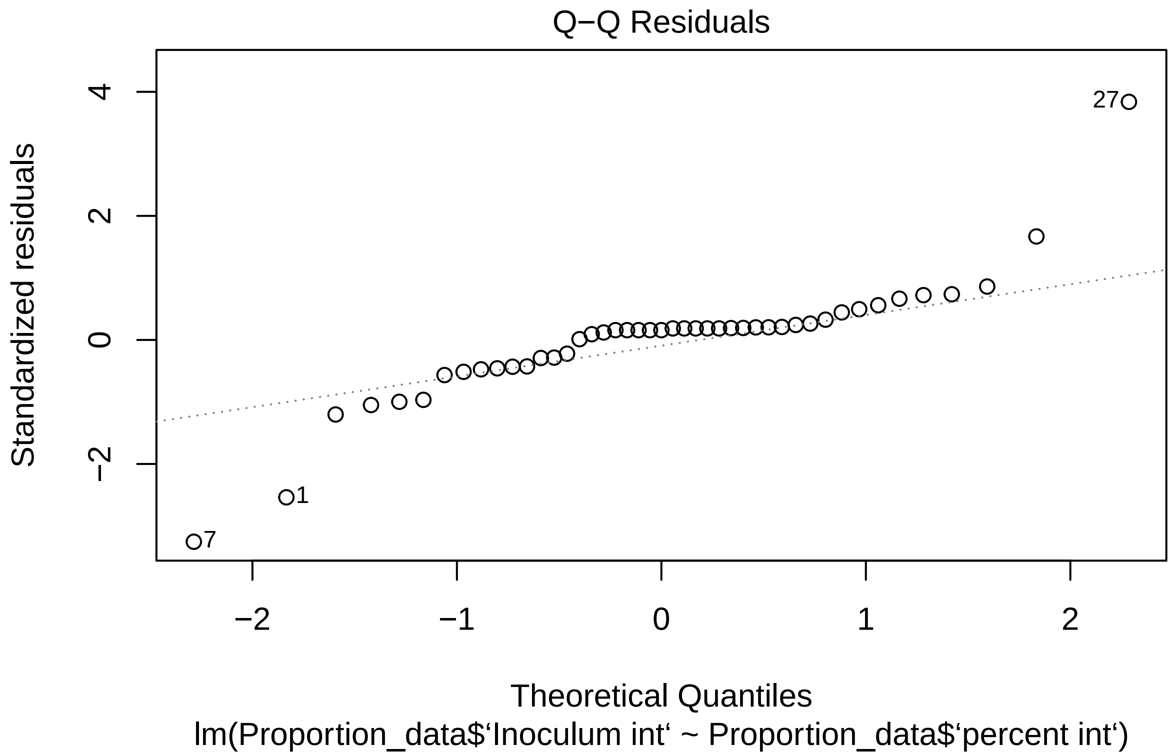
Q−Q Residuals
Standardized residuals
4
2
0
−2
27
1
7
−2
−1
0
1
2
Theoretical Quantiles
lm(Proportion_data$‘Inoculum int‘ ~ Proportion_data$‘percent int‘)

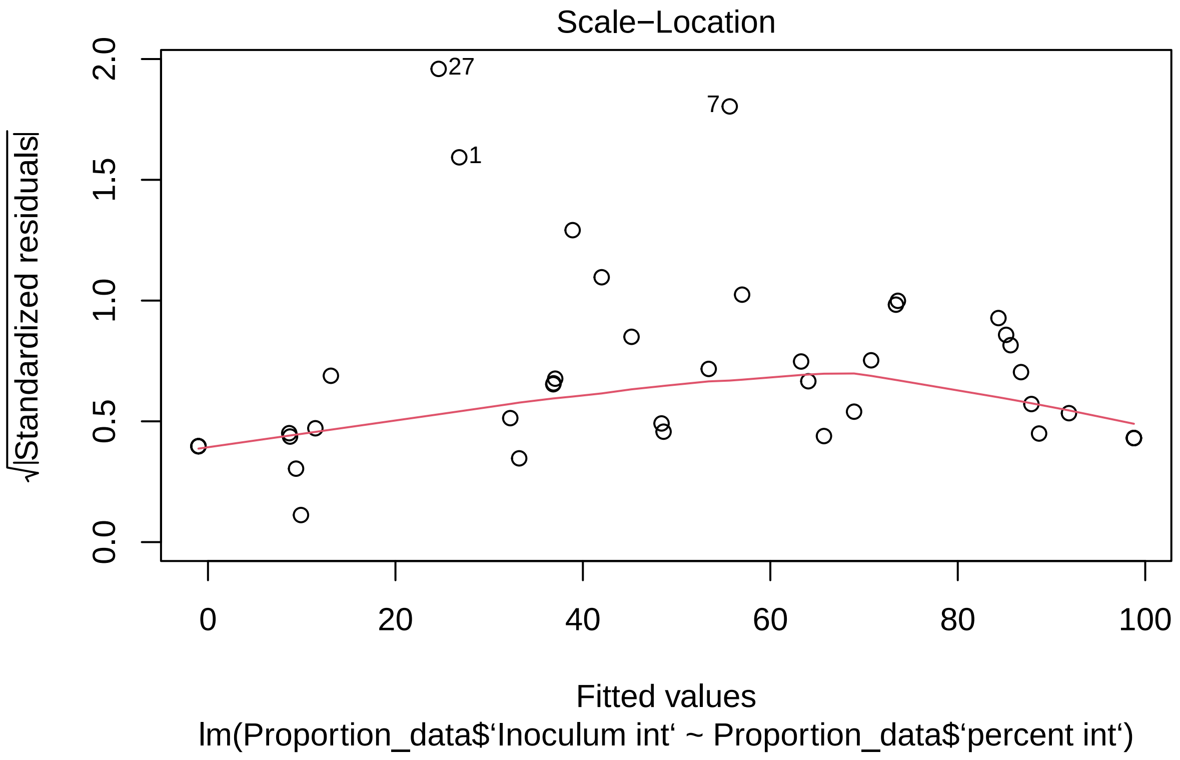
Scale−Location
√|Standardized residuals|
0.0
0.5
1.0
1.5
2.0
27
7
1
0
20
40
60
80
100
Fitted values
lm(Proportion_data$‘Inoculum int‘ ~ Proportion_data$‘percent int‘)

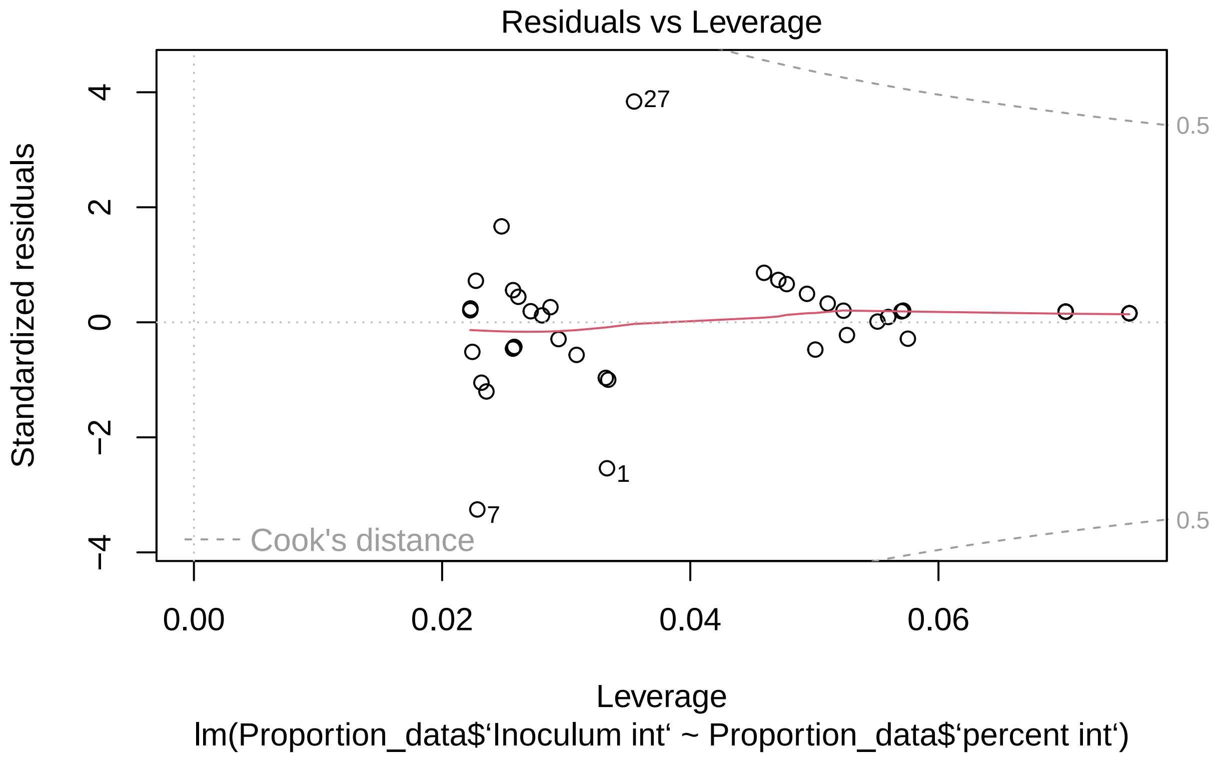


```
plot(proportion_model_size_int)
```

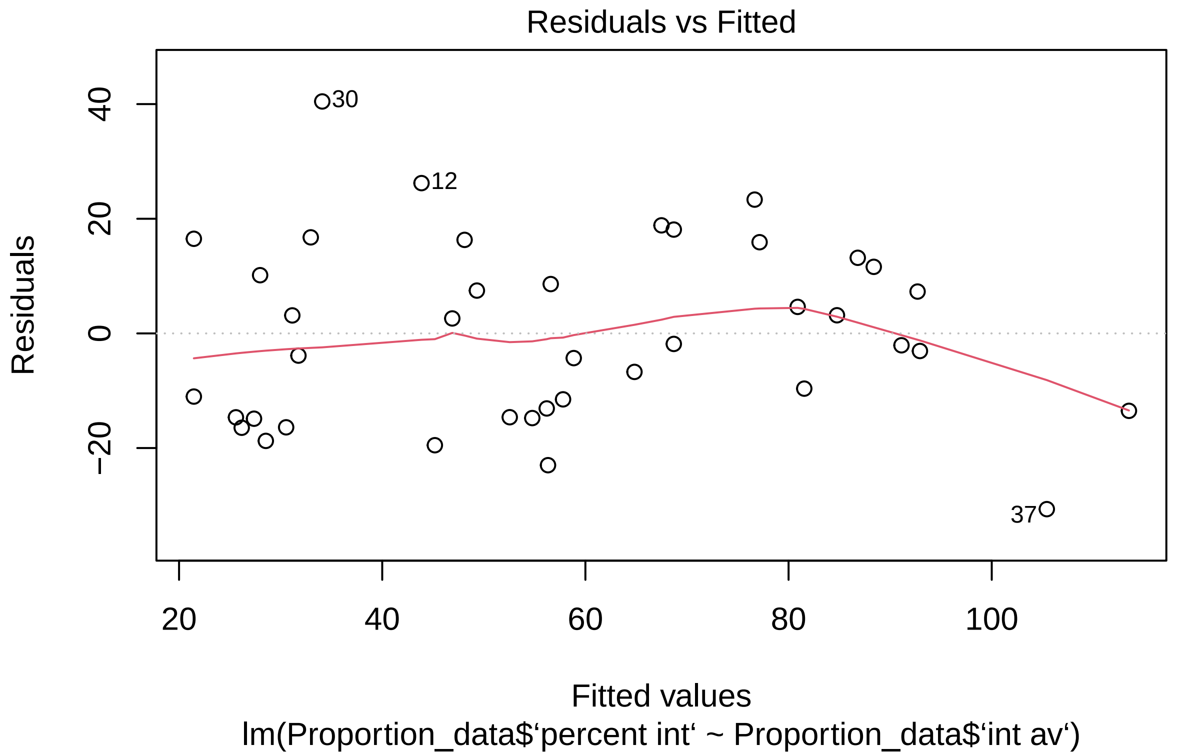

Residuals vs Fitted
Residuals
40
20
0
−20
30
12
37
20
40
60
80
100
Fitted values
lm(Proportion_data$‘percent int‘ ~ Proportion_data$‘int av‘)

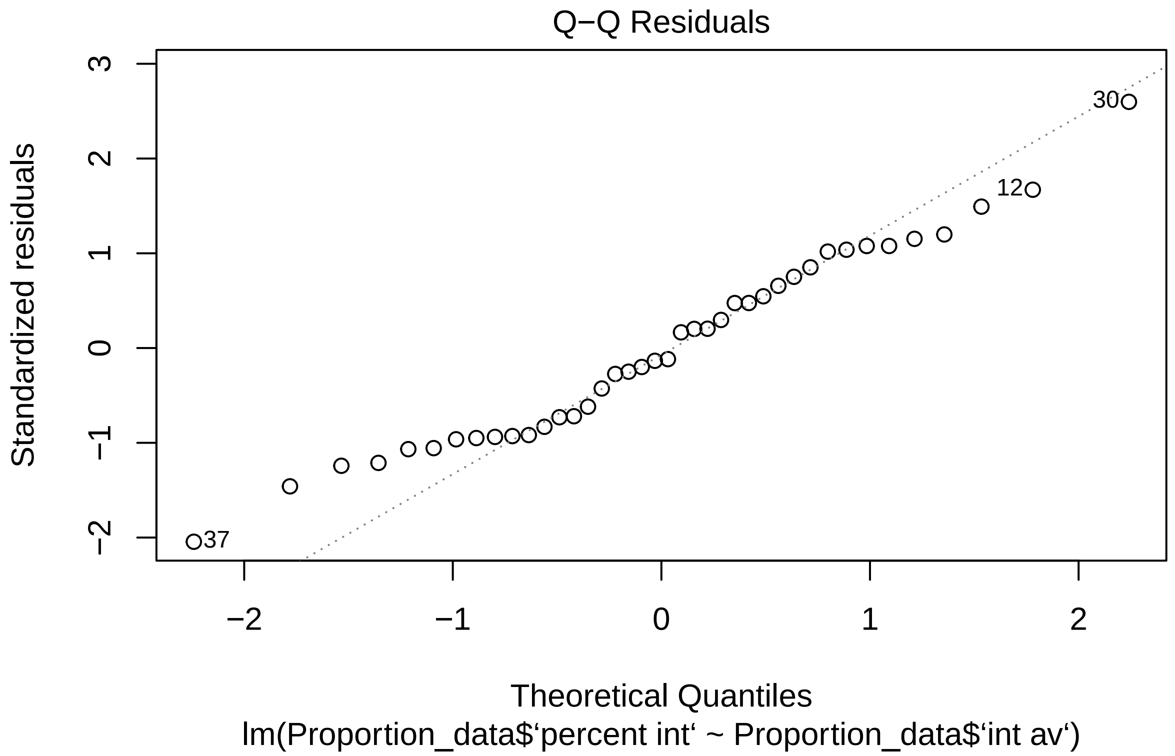
Q−Q Residuals
Standardized residuals
Theoretical Quantiles
lm(Proportion_data$‘percent int‘ ~ Proportion_data$‘int av‘)
3
2
1
0
−1
−2
−2
−1
0
1
2
30
12
37

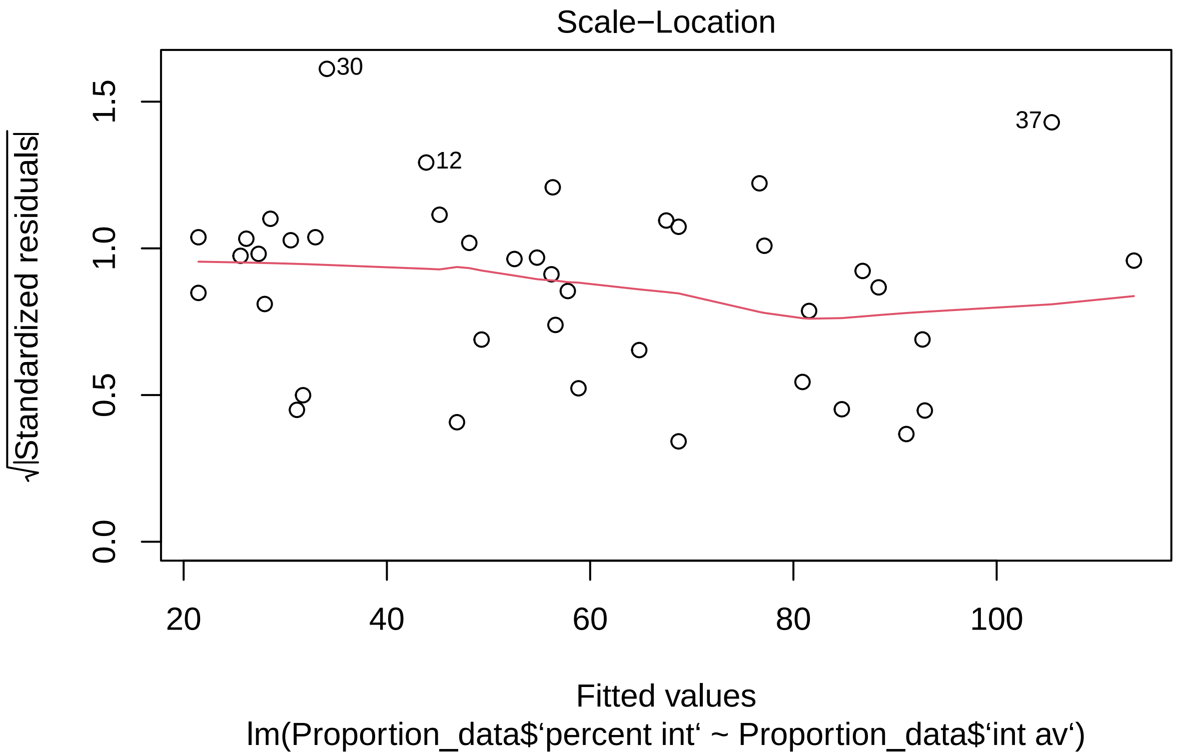
Scale−Location
√|Standardized residuals|
0.0
0.5
1.0
1.5
30
12
37
20
40
60
80
100
Fitted values
lm(Proportion_data$‘percent int‘ ~ Proportion_data$‘int av‘)

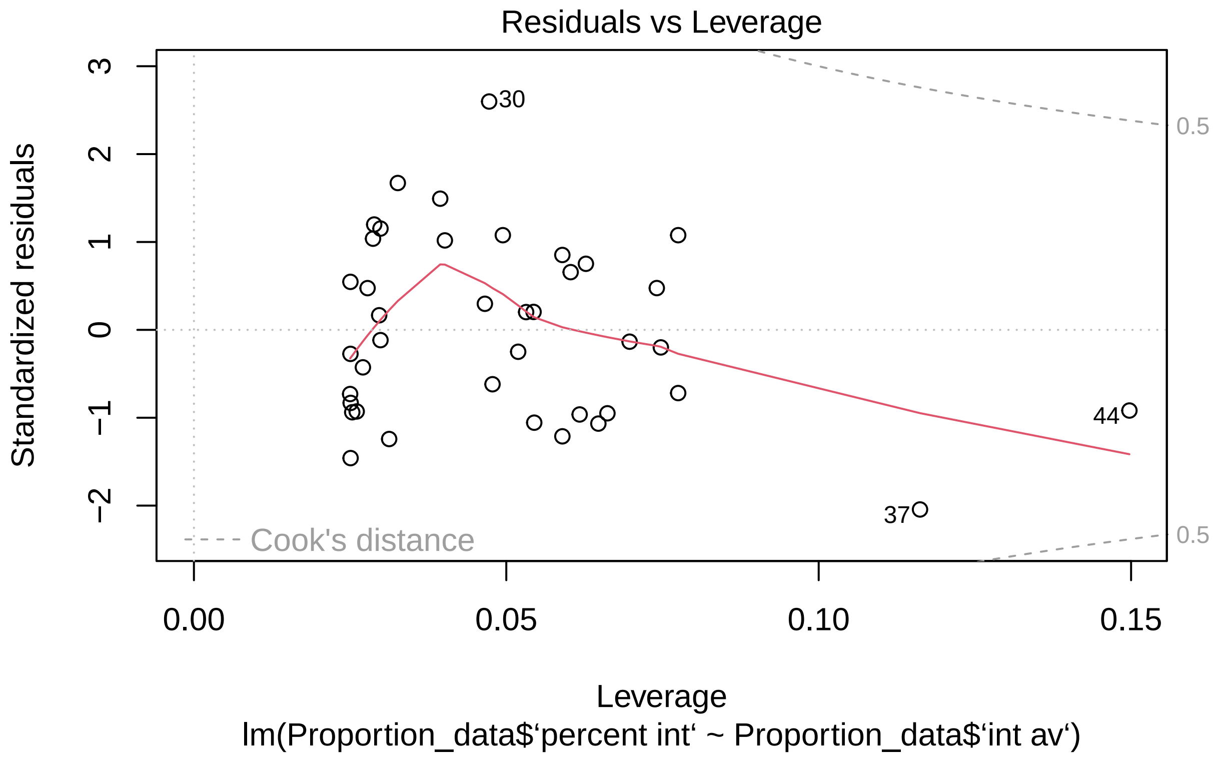


```
plot(proportion_model_size_plus)
```

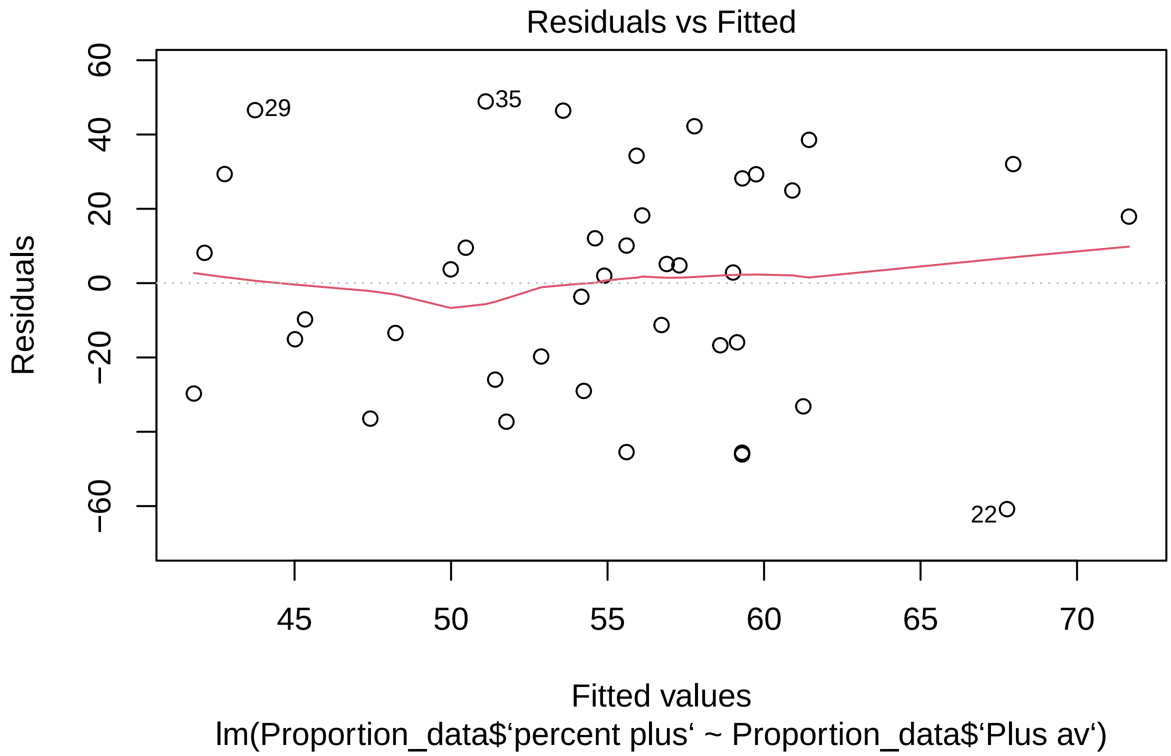
Residuals vs Fitted
Residuals
60
40
20
0
−20
−60
29
35
22
45
50
55
60
65
70
Fitted values
lm(Proportion_data$‘percent plus‘ ~ Proportion_data$‘Plus av‘)

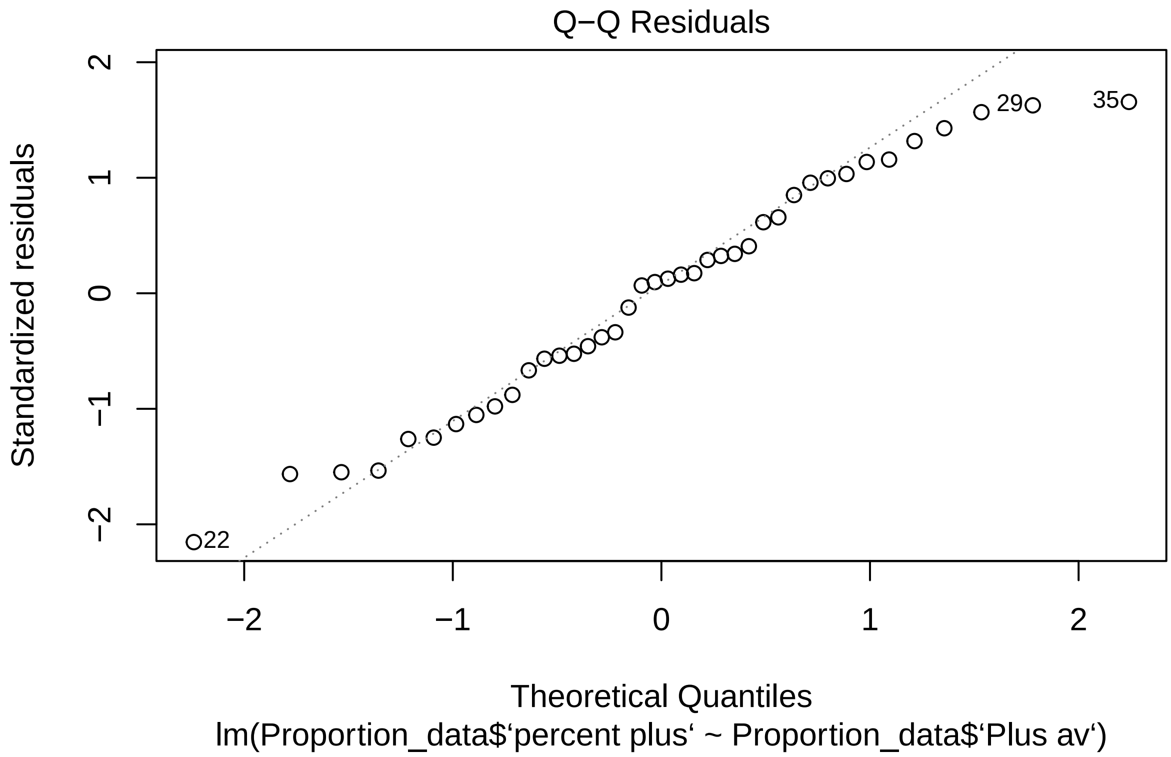
Q−Q Residuals
Standardized residuals
Theoretical Quantiles
lm(Proportion_data$‘percent plus‘ ~ Proportion_data$‘Plus av‘)
2
1
0
−1
−2
−2
−1
0
1
2
22
29
35

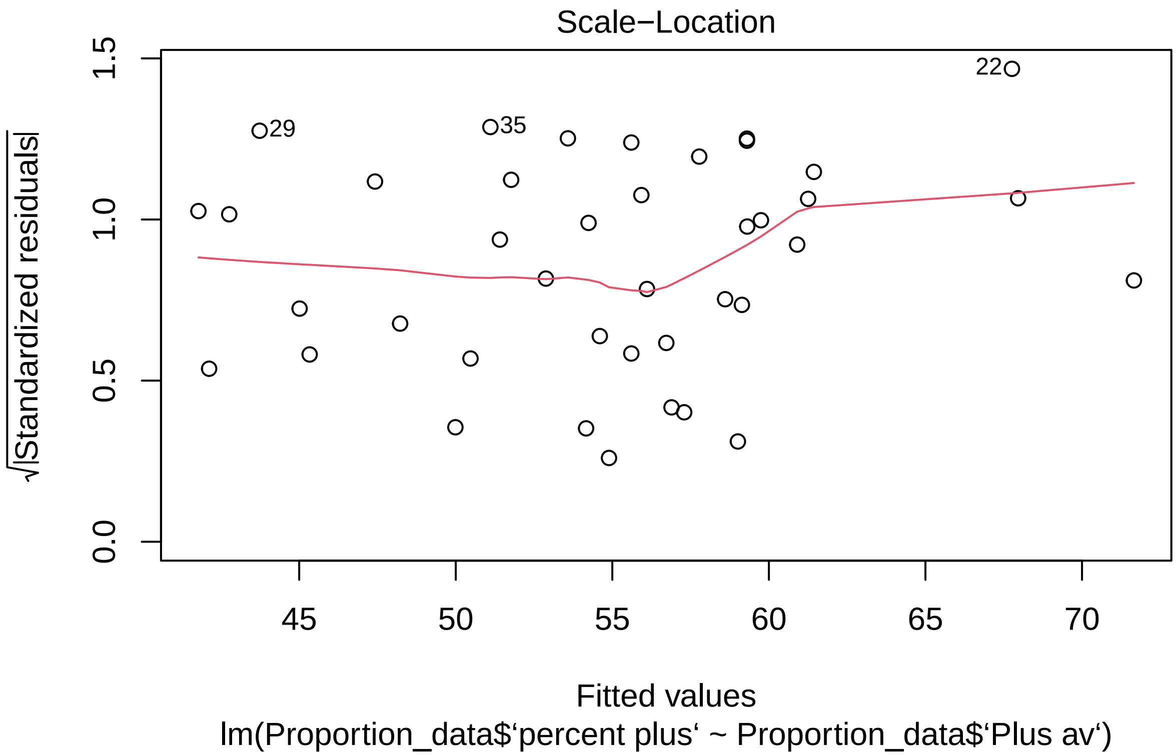
Scale−Location
√|Standardized residuals|
0.0
0.5
1.0
1.5
29
35
22
45
50
55
60
65
70
Fitted values
lm(Proportion_data$‘percent plus‘ ~ Proportion_data$‘Plus av‘)

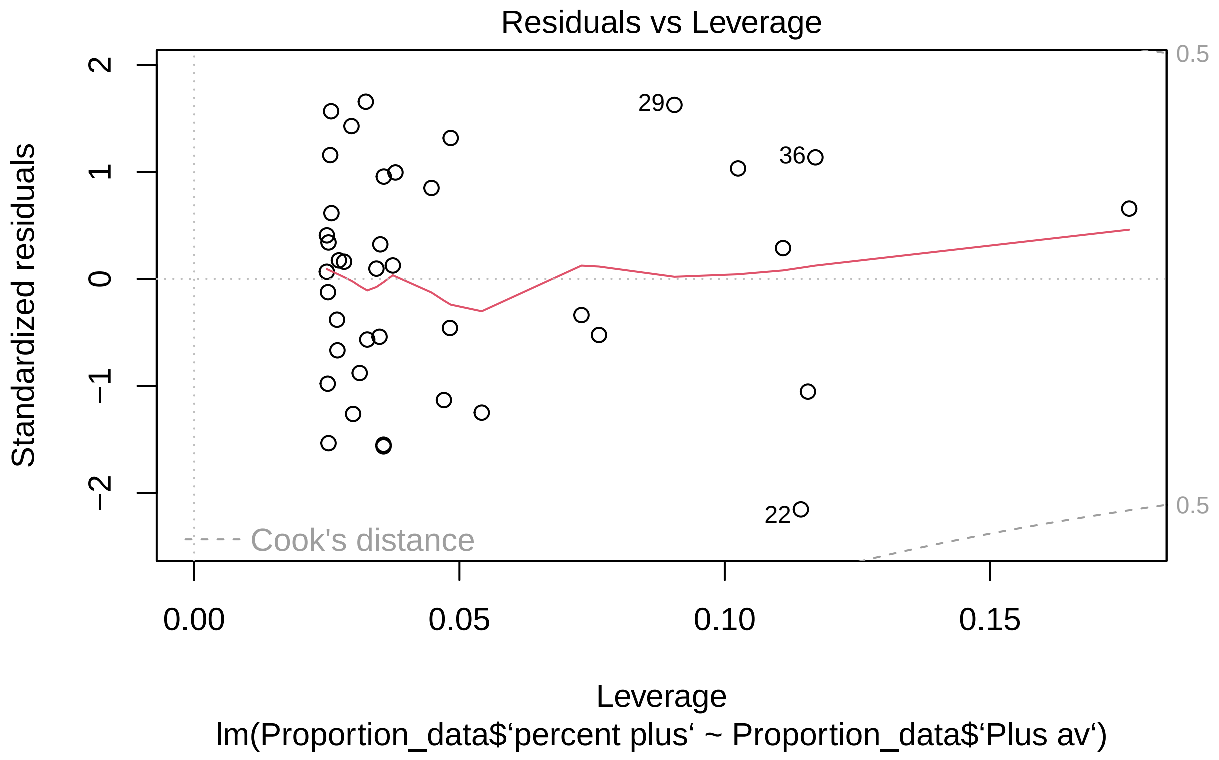


```r
#Here we visually examined the residuals plots for
#each model to check our model assumptions


#We then tested the significance of correlation of these models
summary(proportion_model_inoculum)
```

```
##
## Call:
## lm(formula = Proportion_data$‘Inoculum int‘ ~ Proportion_data$‘percent int‘)
##
## Residuals:
##     Min      1Q  Median      3Q     Max
## -21.665  -2.840   1.021   1.607  25.392
##
## Coefficients:
##                               Estimate Std. Error t value Pr(>|t|)
## (Intercept)                   -1.02078    1.84933  -0.552    0.584
## Proportion_data$‘percent int‘  0.99816    0.03029  32.953   <2e-16 ***
## ---
## Signif. codes:  0 ’***’ 0.001 ’**’ 0.01 ’*’ 0.05 ’.’ 0.1 ’ ’ 1
##
## Residual standard error: 6.735 on 43 degrees of freedom
## Multiple R-squared:  0.9619, Adjusted R-squared:  0.961
## F-statistic:  1086 on 1 and 43 DF,  p-value: < 2.2e-16
```

```r
summary(proportion_model_size_int)
```

```
##
## Call:
## lm(formula = Proportion_data$`percent int` ~ Proportion_data$`int av`)
##
## Residuals:
##     Min      1Q  Median      3Q     Max
## -30.655 -13.787  -1.956  12.005  40.460
##
## Coefficients:
##                            Estimate Std. Error t value Pr(>|t|)
## (Intercept)                 -16.097      7.862  -2.048   0.0476 *
## Proportion_data$`int av`     30.286      3.057   9.908 4.41e-12 ***
## ---
## Signif. codes:  0 '***' 0.001 '**' 0.01 '*' 0.05 '.' 0.1 ' ' 1
##
## Residual standard error: 15.95 on 38 degrees of freedom
##   (5 observations deleted due to missingness)
## Multiple R-squared:  0.721,  Adjusted R-squared:  0.7136
## F-statistic: 98.18 on 1 and 38 DF,  p-value: 4.406e-12
```

```r
summary(proportion_model_size_plus)
```

```
##
## Call:
## lm(formula = Proportion_data$`percent plus` ~ Proportion_data$`Plus av`)
##
## Residuals:
##     Min      1Q  Median      3Q     Max
## -60.818 -21.298   3.285  25.751  48.894
##
## Coefficients:
##                             Estimate Std. Error t value Pr(>|t|)
## (Intercept)                   90.118     24.917   3.617 0.000864 ***
## Proportion_data$`Plus av`     -8.026      5.562  -1.443 0.157214
## ---
## Signif. codes:  0 '***' 0.001 '**' 0.01 '*' 0.05 '.' 0.1 ' ' 1
##
## Residual standard error: 30.01 on 38 degrees of freedom
##   (5 observations deleted due to missingness)
## Multiple R-squared:  0.05195,    Adjusted R-squared:  0.027
## F-statistic: 2.082 on 1 and 38 DF,  p-value: 0.1572
```

```r
#There was a significant positive correlation for our
#models comparing number of nodules with inoculum
#As well as for our model of Fix int nodule size with proportion of nodules
#But not for our model of Fix plus nodule size and proportion of nodules
```

Having shown that the proportion of cheaters impacts the severity of host sanctions we sort to test the effect of spatial seperation using a split root system

```r
experiment <- seperate_split_data$`experiment #`
plant <- seperate_split_data$plant

#These two factors are re named for ease of modelling

seperate_split_size <-
  lmer(seperate_split_data$`Size av` ~ seperate_split_data$treatment
       + (1|experiment) + (1|experiment:plant))
#This nested mixed effects model compares size of nodule
#with the nodule occupant in a pairwise manner
plot(seperate_split_size)
```

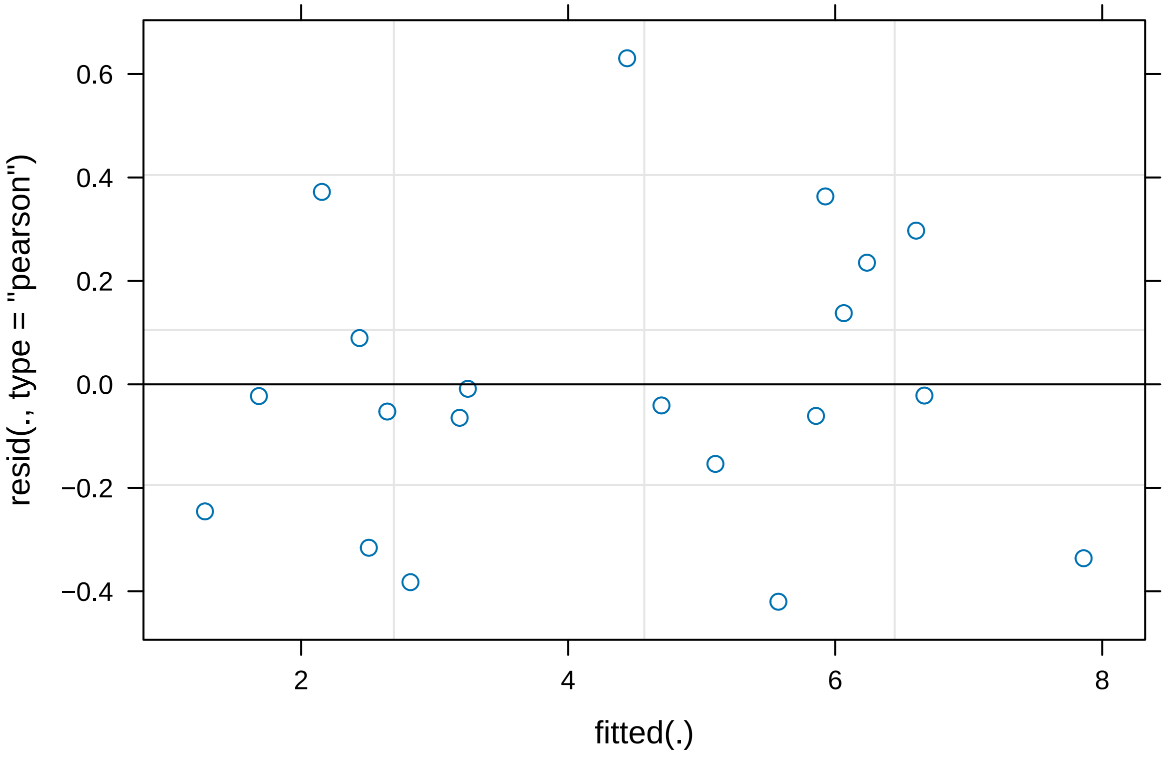


```r
#Plot assessed to test assumptions
summary(seperate_split_size)
```

```
## Linear mixed model fit by REML. t-tests use Satterthwaite's method [
## lmerModLmerTest]
## Formula: seperate_split_data$'Size av' ~ seperate_split_data$treatment +
##     (1 | experiment) + (1 | experiment:plant)
##
## REML criterion at convergence: 42
##
## Scaled residuals:
##      Min       1Q   Median       3Q      Max
## -1.09428 -0.46023 -0.08253  0.42212  1.64195
```

```
##
## Random effects:
##  Groups           Name        Variance Std.Dev.
##  experiment:plant (Intercept) 0.5420   0.7362
##  experiment       (Intercept) 0.6474   0.8046
##  Residual                     0.1475   0.3840
## Number of obs: 20, groups:  experiment:plant, 10; experiment, 2
##
## Fixed effects:
##                                   Estimate Std. Error     df t value Pr(>|t|)
## (Intercept)                         2.8465     0.6342 1.0153   4.488    0.137
## seperate_split_data$treatmentPlus   3.4191     0.1717 9.0000  19.909 9.45e-09
##
## (Intercept)
## seperate_split_data$treatmentPlus ***
## ---
## Signif. codes:  0 '***' 0.001 '**' 0.01 '*' 0.05 '.' 0.1 ' ' 1
##
## Correlation of Fixed Effects:
##             (Intr)
## sprt_spl_$P -0.135
```

```
#There was a signficant differnce in size of nodule based on occupant

split_proportion <-
  lm(seperate_split_data$Percentage ~ seperate_split_data$`size int`)
#This model tests the correlation between
#proportion of nodules occupied by fix int and their size

plot(split_proportion)
```

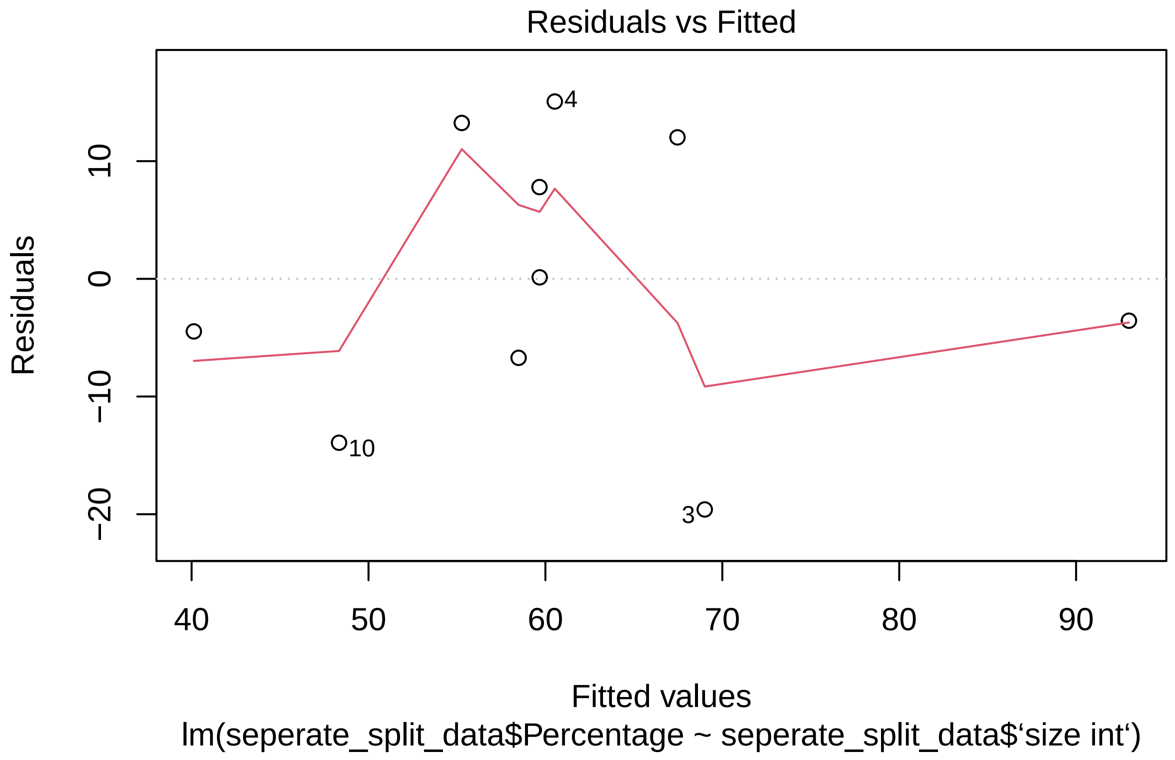
Residuals vs Fitted
Residuals
10
0
−10
−20
4
10
3
40
50
60
70
80
90
Fitted values
lm(seperate_split_data$Percentage ~ seperate_split_data$‘size int‘)

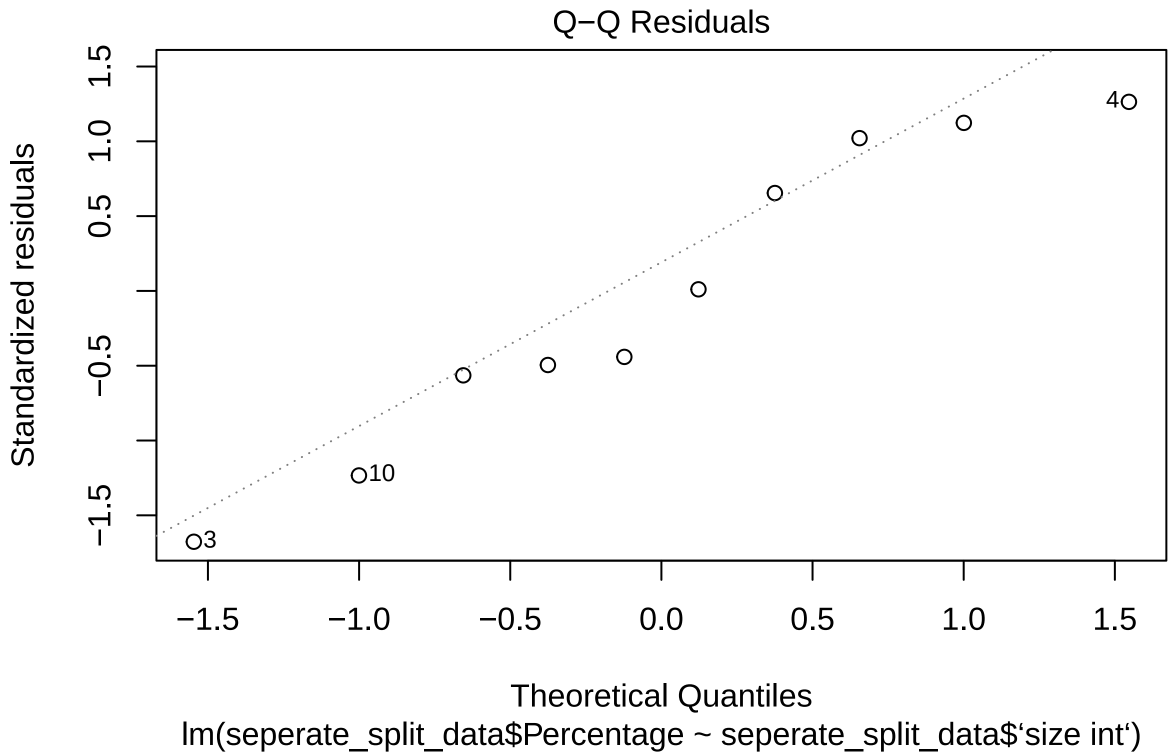
Q−Q Residuals
Standardized residuals
1.5
1.0
0.5
−0.5
−1.5
4
10
3
−1.5
−1.0
−0.5
0.0
0.5
1.0
1.5
Theoretical Quantiles
lm(seperate_split_data$Percentage ~ seperate_split_data$‘size int‘)

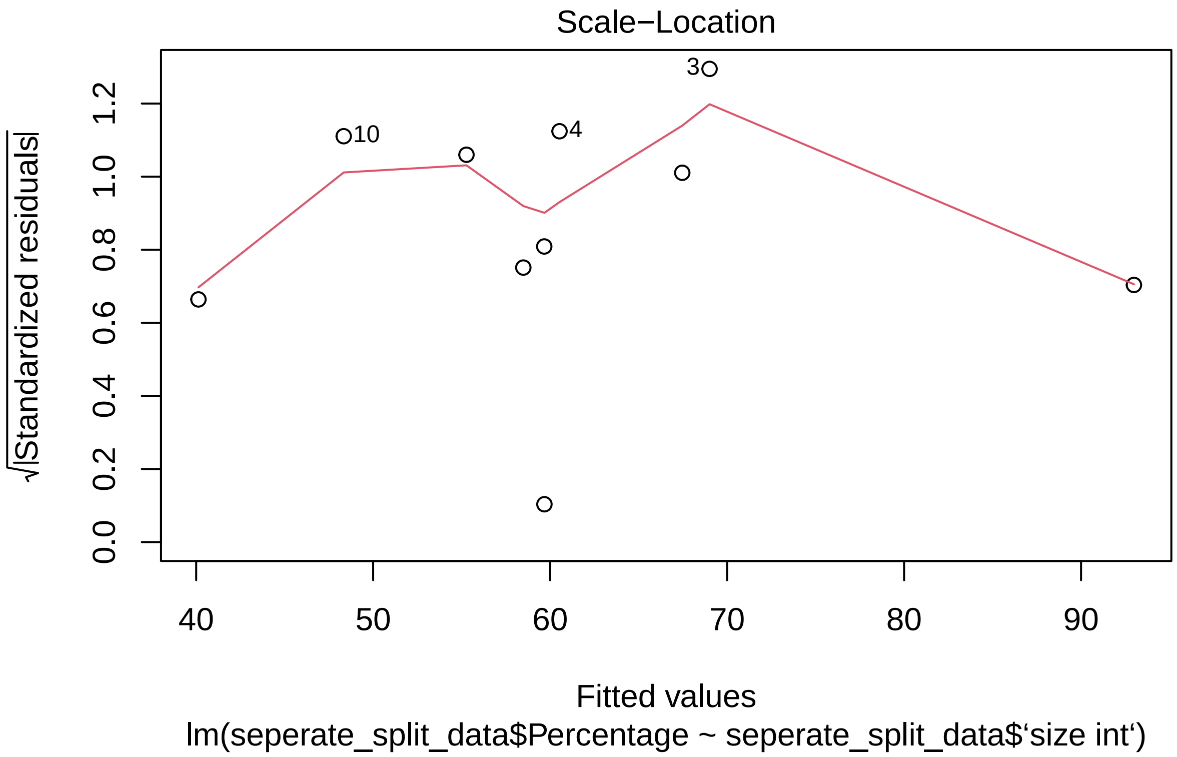
Scale−Location
√|Standardized residuals|
0.0
0.2
0.4
0.6
0.8
1.0
1.2
40
50
60
70
80
90
10
4
3
Fitted values
lm(seperate_split_data$Percentage ~ seperate_split_data$‘size int‘)

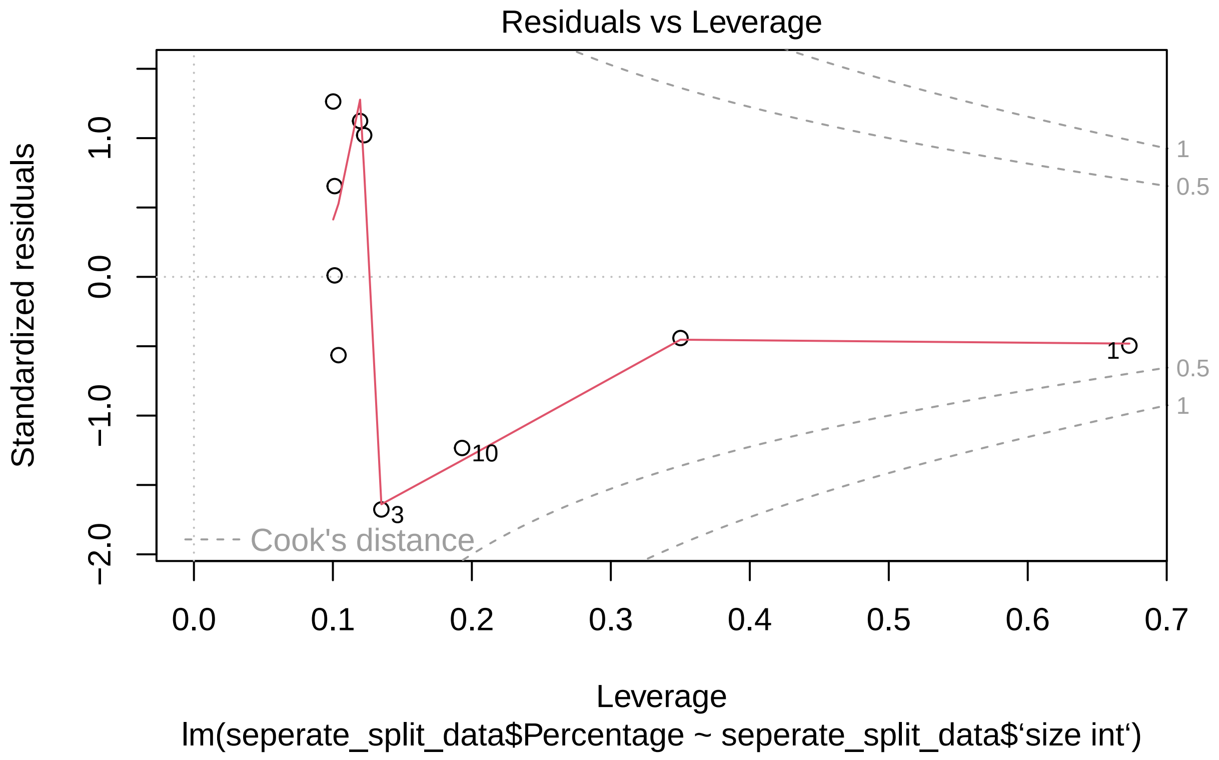


```r
#plot for model assumptions

summary(split_proportion)
```

```
##
## Call:
## lm(formula = seperate_split_data$Percentage ~ seperate_split_data$‘size int‘)
##
## Residuals:
##     Min      1Q  Median      3Q     Max
## -19.596  -6.151  -1.715  10.965  15.066
##
## Coefficients:
##                                Estimate Std. Error t value Pr(>|t|)
## (Intercept)                      26.581     11.070   2.401   0.0431 *
## seperate_split_data$‘size int‘   13.091      3.912   3.346   0.0101 *
## ---
## Signif. codes:  0 ’***’ 0.001 ’**’ 0.01 ’*’ 0.05 ’.’ 0.1 ’ ’ 1
##
## Residual standard error: 12.57 on 8 degrees of freedom
##   (10 observations deleted due to missingness)
## Multiple R-squared:  0.5832, Adjusted R-squared:  0.5312
## F-statistic:  11.2 on 1 and 8 DF,  p-value: 0.01014
```

```r
#There is a significant positive correlation
#between proportion of nodules occupied and fix int nodule size
```

We next sort to test for effects caused by the split root system itself so repeated all of the above steps for split root plants with mixed inocula

```r
experiment <- as.factor(mixed_split_data_plant$Experiment)
plant <- as.factor(mixed_split_data_plant$plant)

mixed_plant_size <-
  lmer(mixed_split_data_plant$`Size av` ~ mixed_split_data_plant$treatment
       + (1|experiment) + (1|experiment/plant))

plot(mixed_plant_size)
```

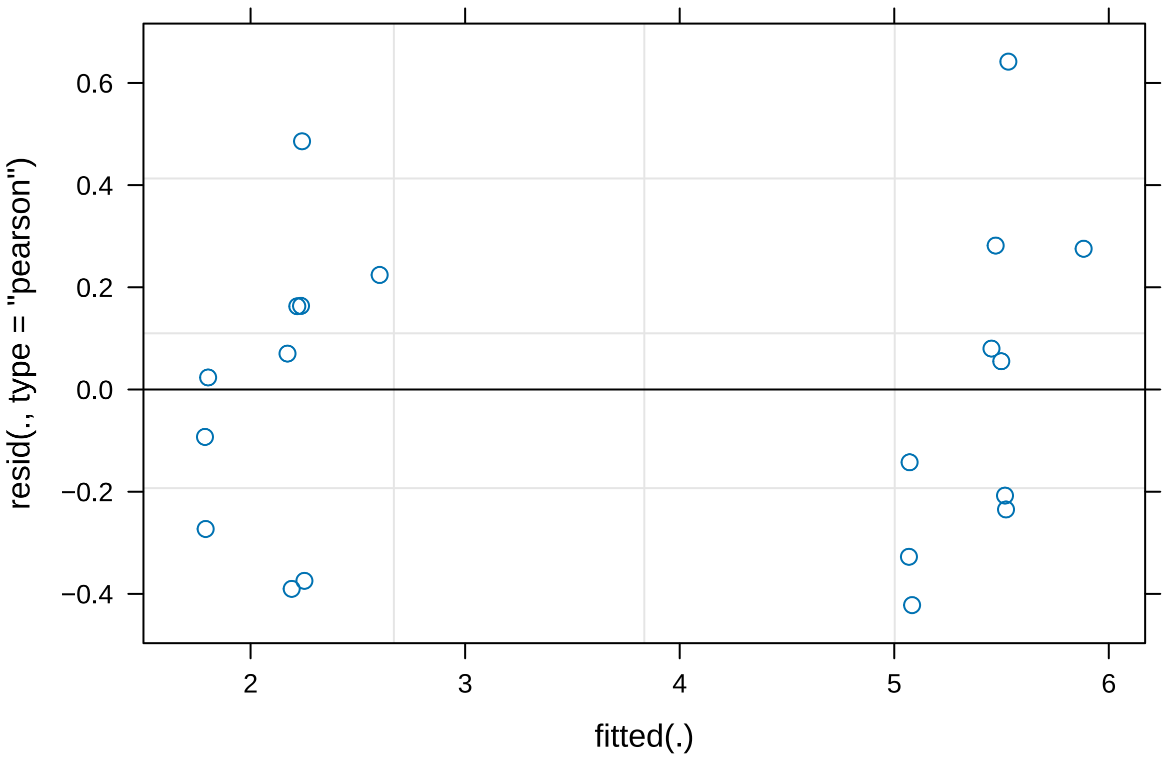


```r
summary(mixed_plant_size)
```

```
## Linear mixed model fit by REML. t-tests use Satterthwaite's method [
## lmerModLmerTest]
## Formula: mixed_split_data_plant$'Size av' ~ mixed_split_data_plant$treatment +
##     (1 | experiment) + (1 | experiment/plant)
##
## REML criterion at convergence: 28
##
```

```
## Scaled residuals:
##     Min      1Q  Median      3Q     Max
## -1.1494 -0.6655  0.1074  0.4871  1.7479
##
## Random effects:
##  Groups           Name        Variance Std.Dev.
##  plant.experiment (Intercept) 0.09088  0.3015
##  experiment       (Intercept) 0.01557  0.1248
##  experiment.1     (Intercept) 0.02212  0.1487
##  Residual                     0.13485  0.3672
## Number of obs: 20, groups:  plant:experiment, 10; experiment, 2
##
## Fixed effects:
##                                        Estimate Std. Error     df t value
## (Intercept)                              2.1678     0.2072 1.1944   10.46
## mixed_split_data_plant$treatmentPlus   3.2811     0.1642 8.9999   19.98
##                                        Pr(>|t|)
## (Intercept)                              0.0403 *
## mixed_split_data_plant$treatmentPlus 9.17e-09 ***
## ---
## Signif. codes:  0 '***' 0.001 '**' 0.01 '*' 0.05 '.' 0.1 ' ' 1
##
## Correlation of Fixed Effects:
##             (Intr)
## mxd_spl__$P -0.396
```

```
mixed_proportion <-
  lm(mixed_split_data_plant$Percentage~mixed_split_data_plant$`size int`)

plot(mixed_proportion)
```

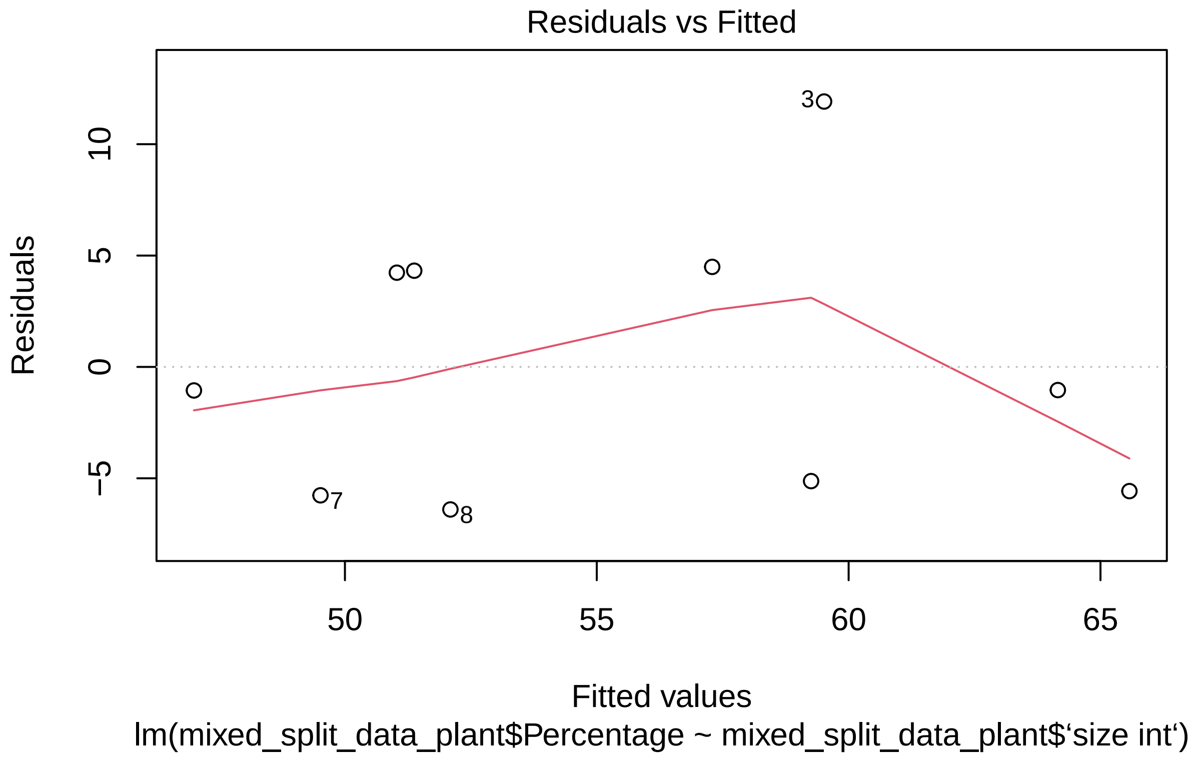
Residuals vs Fitted
Residuals
Fitted values
lm(mixed_split_data_plant$Percentage ~ mixed_split_data_plant$‘size int‘)
10
5
0
−5
50
55
60
65
3
7
8

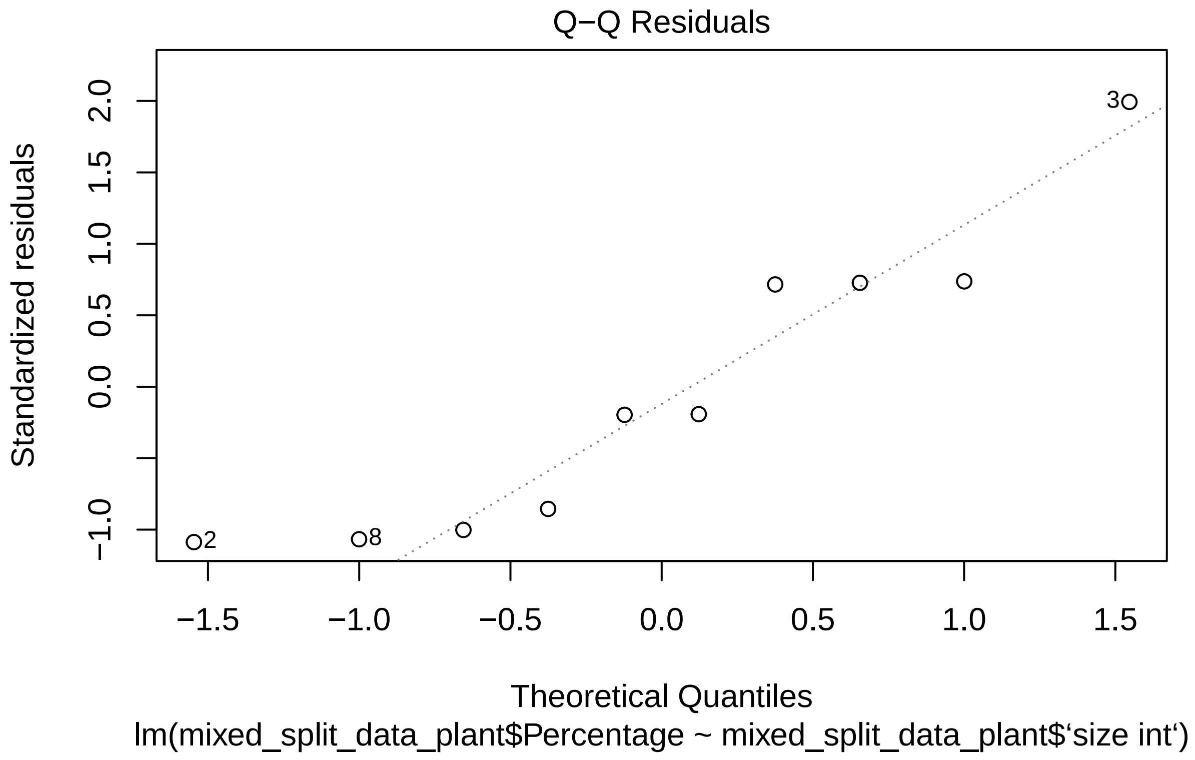
Q−Q Residuals
Standardized residuals
2.0
1.5
1.0
0.5
0.0
−1.0
3
2
8
−1.5
−1.0
−0.5
0.0
0.5
1.0
1.5
Theoretical Quantiles
lm(mixed_split_data_plant$Percentage ~ mixed_split_data_plant$‘size int‘)

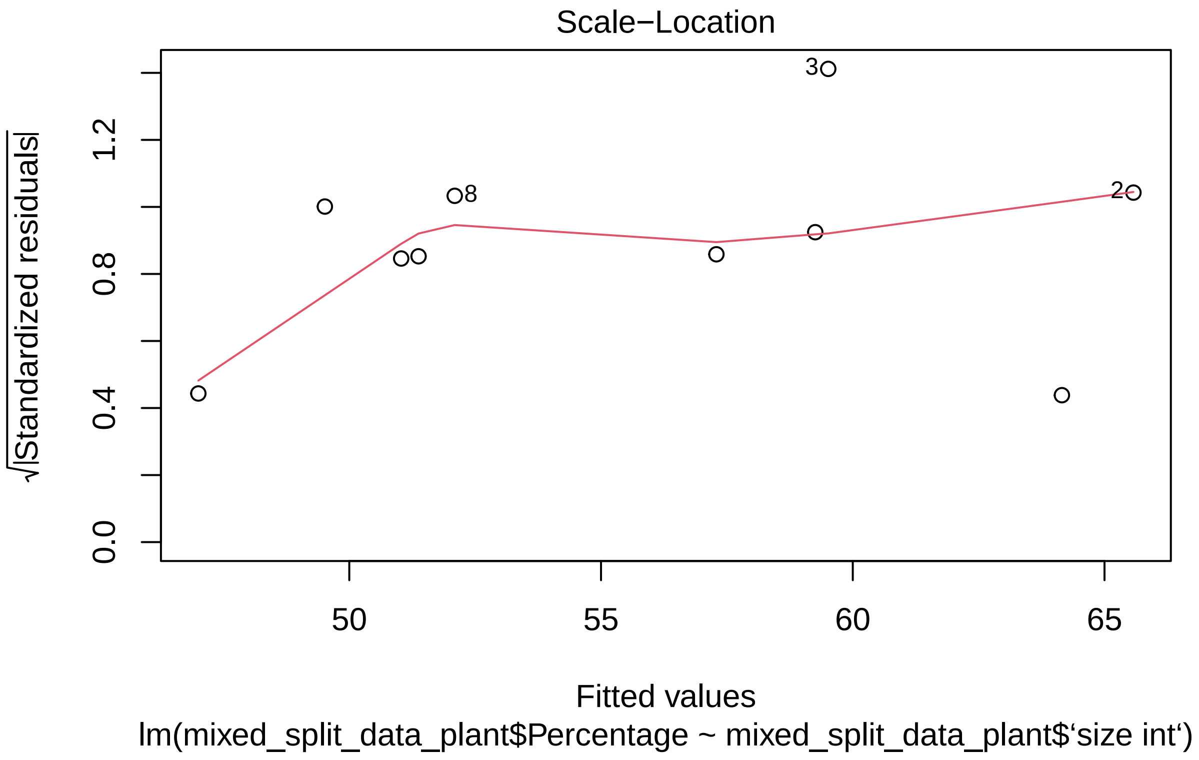
Scale−Location
√|Standardized residuals|
0.0
0.4
0.8
1.2
50
55
60
65
3
8
2
Fitted values
lm(mixed_split_data_plant$Percentage ~ mixed_split_data_plant$‘size int‘)

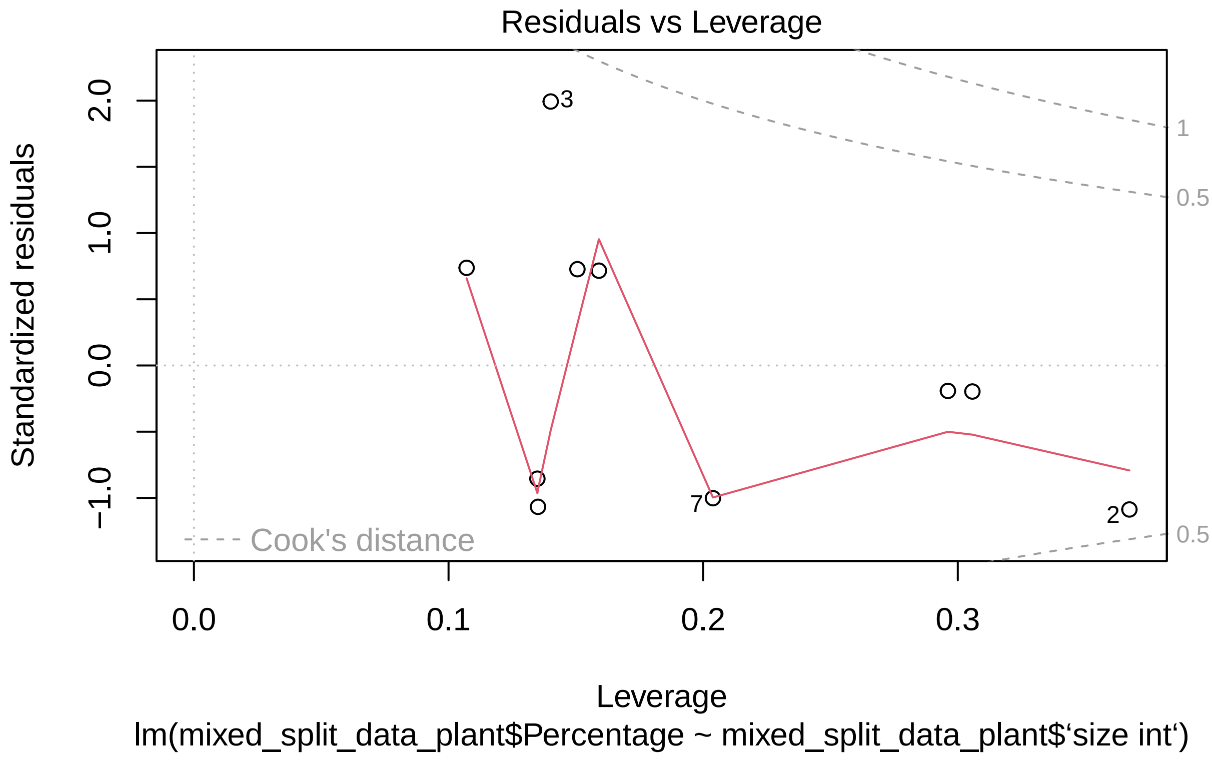


```
summary(mixed_proportion)
```

```
##
## Call:
## lm(formula = mixed_split_data_plant$Percentage ~ mixed_split_data_plant$‘size int‘)
##
## Residuals:
##    Min     1Q Median     3Q    Max
## -6.399 -5.463 -1.048  4.298 11.915
##
## Coefficients:
##                                   Estimate Std. Error t value Pr(>|t|)
## (Intercept)                         25.458     10.385   2.451   0.0398 *
## mixed_split_data_plant$‘size int‘   14.196      4.783   2.968   0.0179 *
## ---
## Signif. codes:  0 ’***’ 0.001 ’**’ 0.01 ’*’ 0.05 ’.’ 0.1 ’ ’ 1
##
## Residual standard error: 6.447 on 8 degrees of freedom
##   (10 observations deleted due to missingness)
## Multiple R-squared:  0.5241, Adjusted R-squared:  0.4646
## F-statistic: 8.809 on 1 and 8 DF,  p-value: 0.01792
```

The results were the same as in the seperated split roots To confirm the absence of proximity effects we compared the mixed and separate split root results

```r
Size <- mixedvseperate$`Size av`

treatment <- mixedvseperate$treatment

#We first defined the factors for model construction

mixedvseperatemodel <- lm(Size ~ treatment)

#This model compares the int and plus nodules
#for both split root types (mixed and seperate)

plot(mixedvseperatemodel)
```

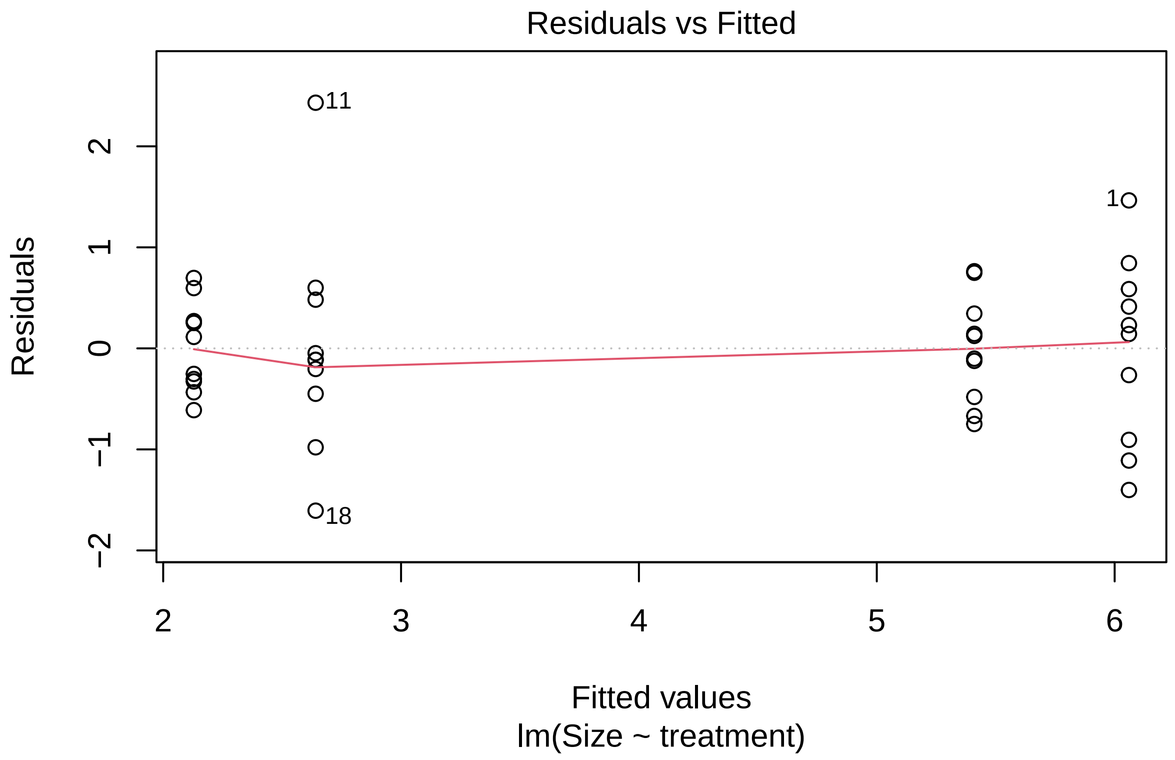

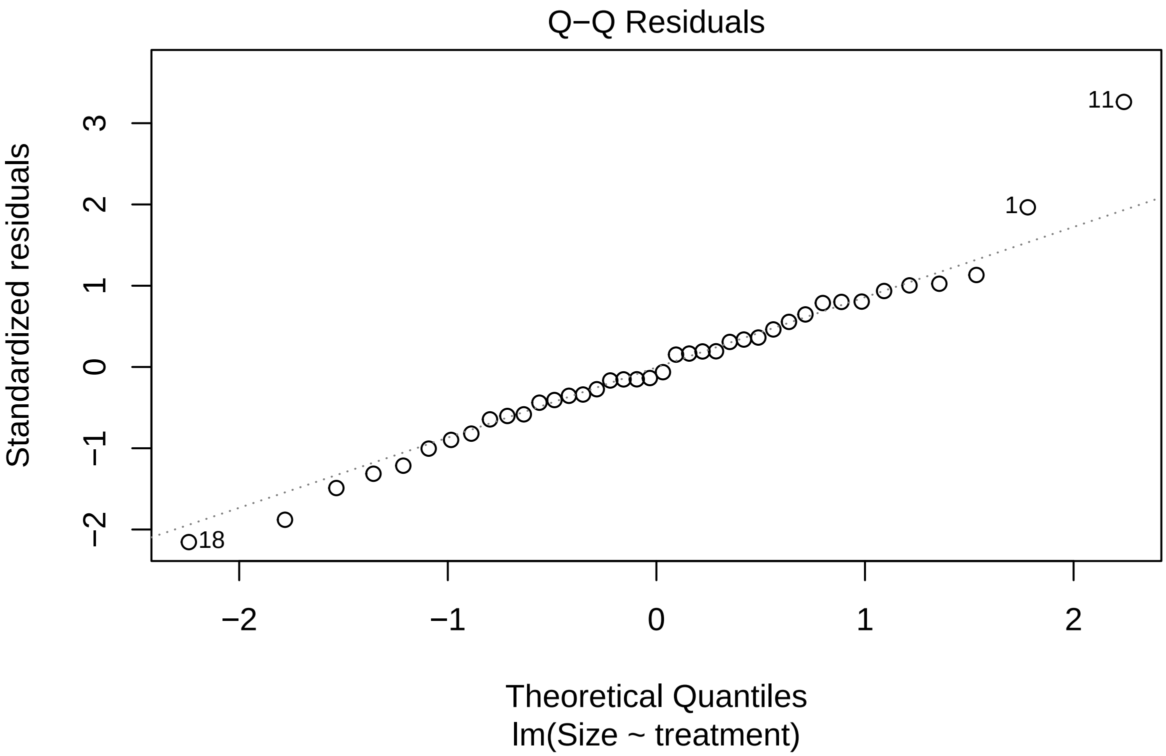

Q−Q Residuals
Standardized residuals
Theoretical Quantiles
lm(Size ~ treatment)
−2
−1
0
1
2
3
11
1
18

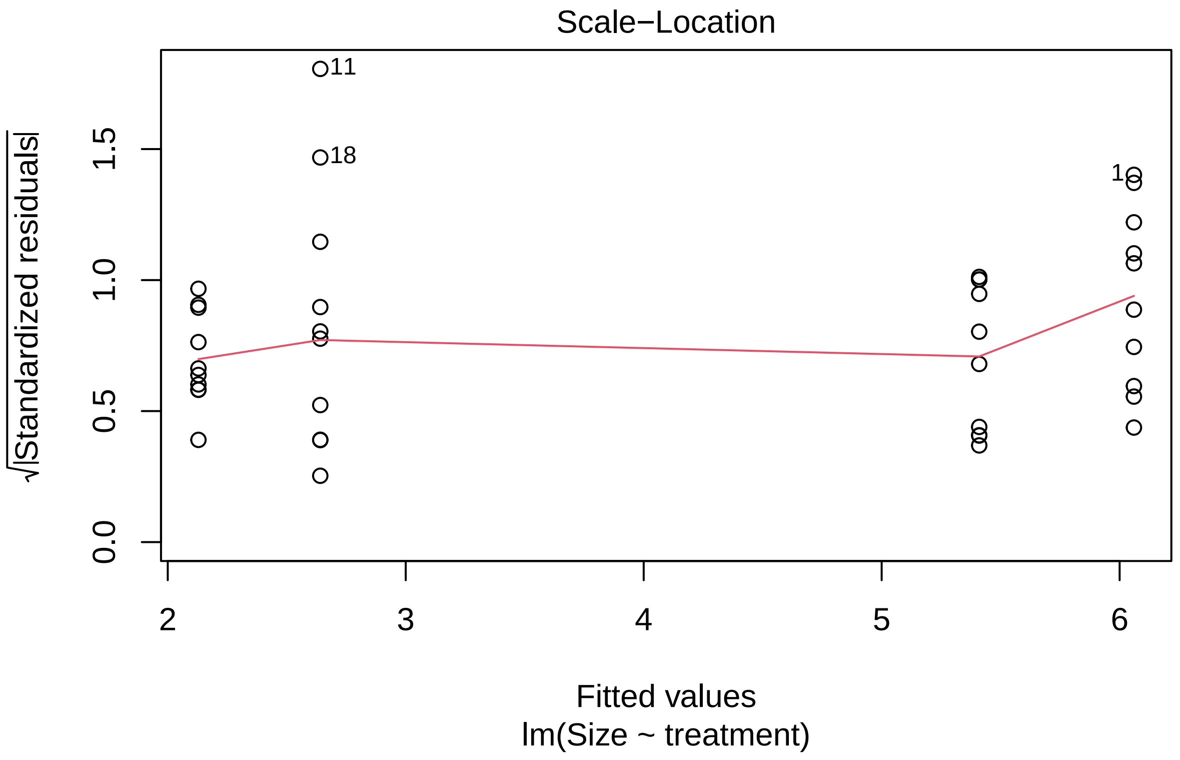
Scale−Location
√|Standardized residuals|
0.0
0.5
1.0
1.5
11
18
2
3
4
5
6
Fitted values
lm(Size ~ treatment)

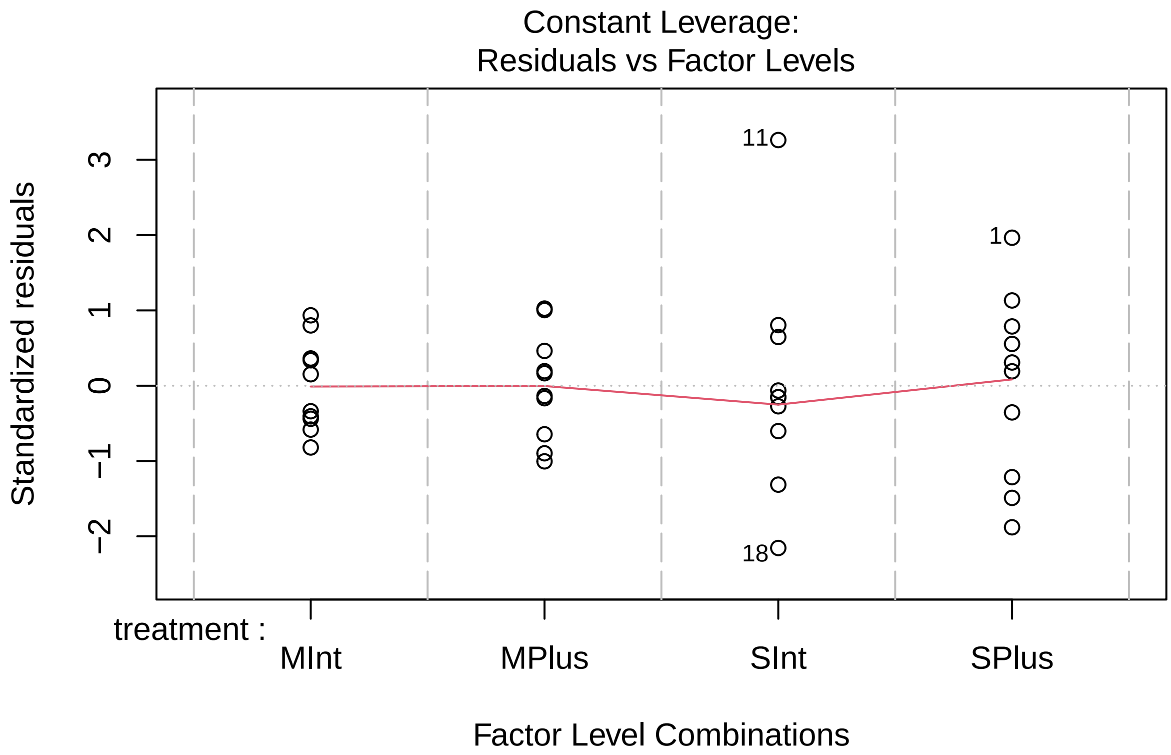


```r
anova(mixedvseperatemodel)
```

```
## Analysis of Variance Table
##
## Response: Size
##           Df  Sum Sq Mean Sq F value    Pr(>F)
## treatment  3 115.656  38.552  62.434 2.434e-14 ***
## Residuals 36  22.229   0.617
## ---
## Signif. codes:  0 '***' 0.001 '**' 0.01 '*' 0.05 '.' 0.1 ' ' 1
```

```r
#this anova showed their were significant differences so
#we carried out a post hoc test to check which
#comparisons were significantly different

MvSanova <- aov(mixedvseperatemodel)

TukeyHSD(MvSanova)
```

```
##   Tukey multiple comparisons of means
##     95% family-wise confidence level
##
## Fit: aov(formula = mixedvseperatemodel)
##
## $treatment
```

```
##                  diff        lwr       upr     p adj
## MPlus-MInt   3.28106  2.3346046  4.227515 0.0000000
## SInt-MInt    0.51203 -0.4344254  1.458485 0.4733113
## SPlus-MInt   3.93117  2.9847146  4.877625 0.0000000
## SInt-MPlus  -2.76903 -3.7154854 -1.822575 0.0000000
## SPlus-MPlus  0.65011 -0.2963454  1.596565 0.2674440
## SPlus-SInt   3.41914  2.4726846  4.365595 0.0000000
```

```
#There was no significant difference between
#nodules of the same type on the different split root systems
```

We concluded that there was no proximity effect of sanctioning and therefore sanctioning is based on a global comparison

We next tested the sanctioning of strains with small variations in fixation effectiveness

```
#We compared the size of nodules for each of the
#three strains when inoculated with either itself or the other two strains

Plus_competition_model <-
  lm(variable_fixers_competition$Plus_average
     ~ variable_fixers_competition$plus_nodule_type)

L_competition_model <-
  lm(variable_fixers_competition$L_average
     ~ variable_fixers_competition$L_nodule_type)

Int_competition_model <-
  lm(variable_fixers_competition$Int_average
     ~ variable_fixers_competition$Int_nodule_type)

#We then tested the assumptions of our models and carried out one way ANOVA's

plot(Plus_competition_model)
```

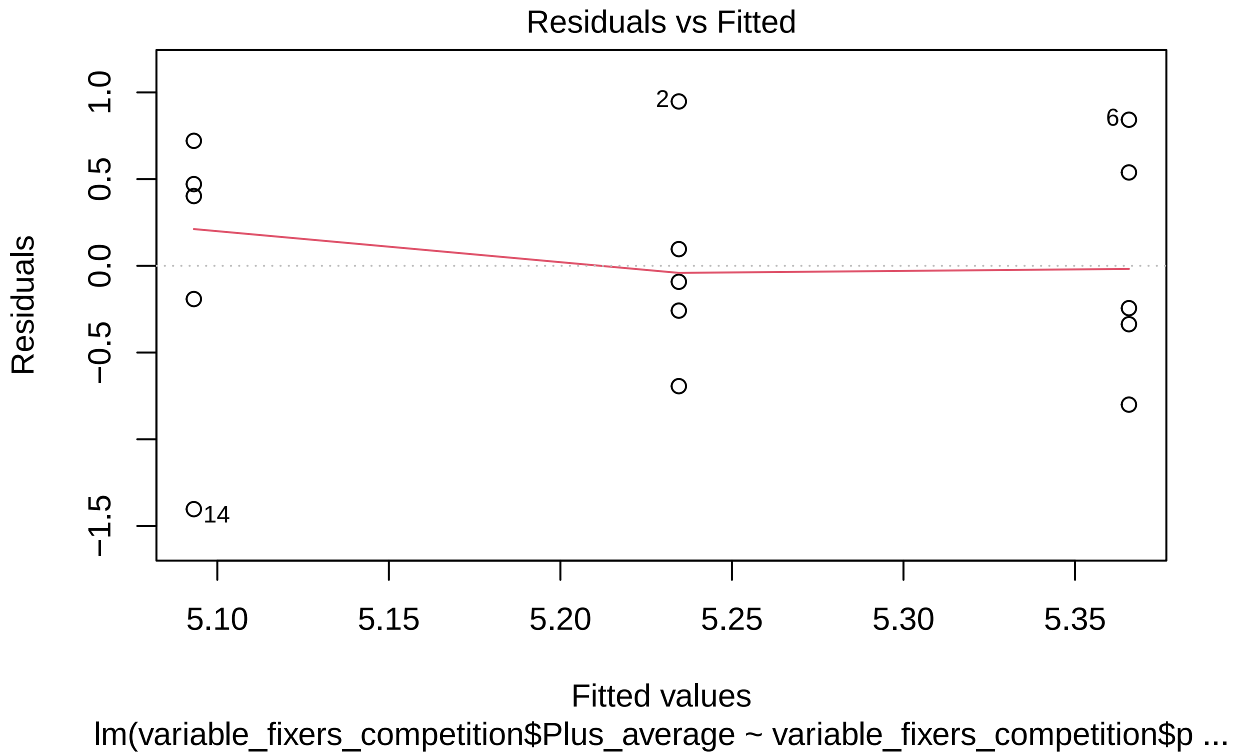
Residuals vs Fitted
Residuals
1.0
0.5
0.0
−0.5
−1.5
2
6
14
5.10
5.15
5.20
5.25
5.30
5.35
Fitted values
lm(variable_fixers_competition$Plus_average ~ variable_fixers_competition$p ...

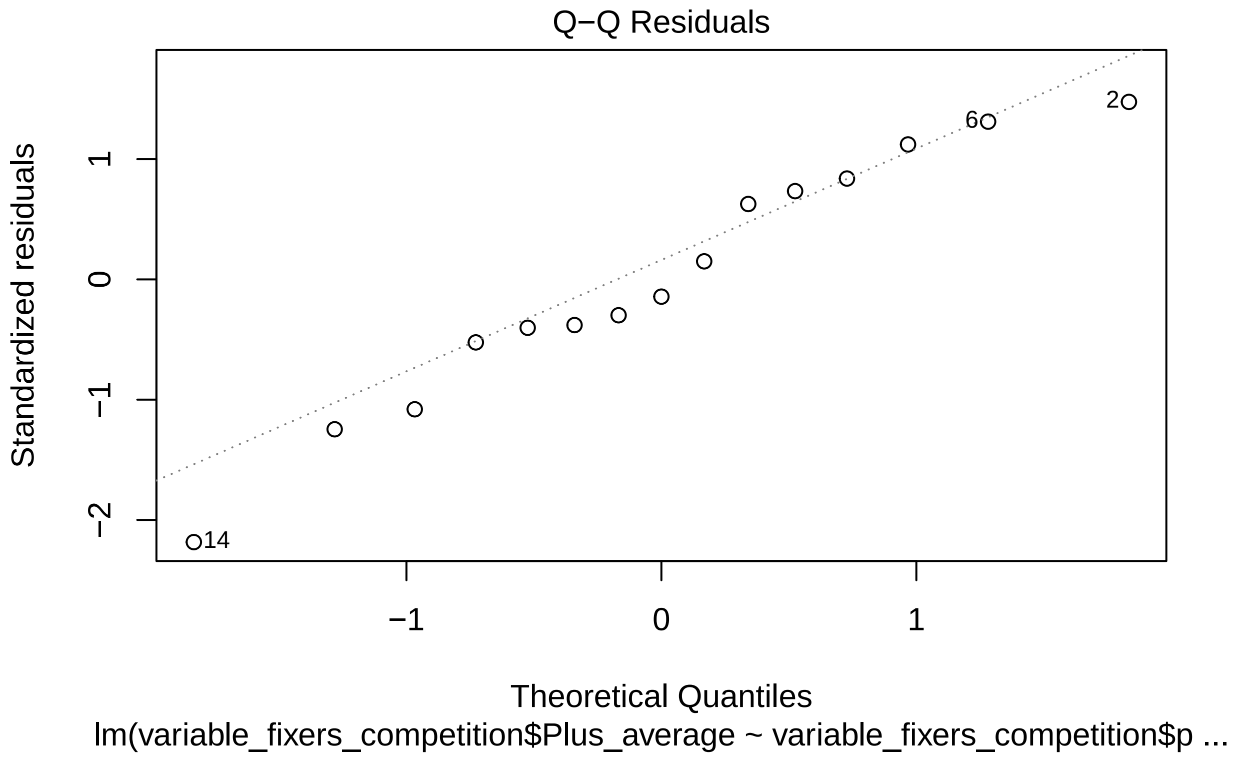
Q−Q Residuals
Standardized residuals
Theoretical Quantiles
lm(variable_fixers_competition$Plus_average ~ variable_fixers_competition$p ...
−2
−1
0
1
−1
0
1
14
6
2

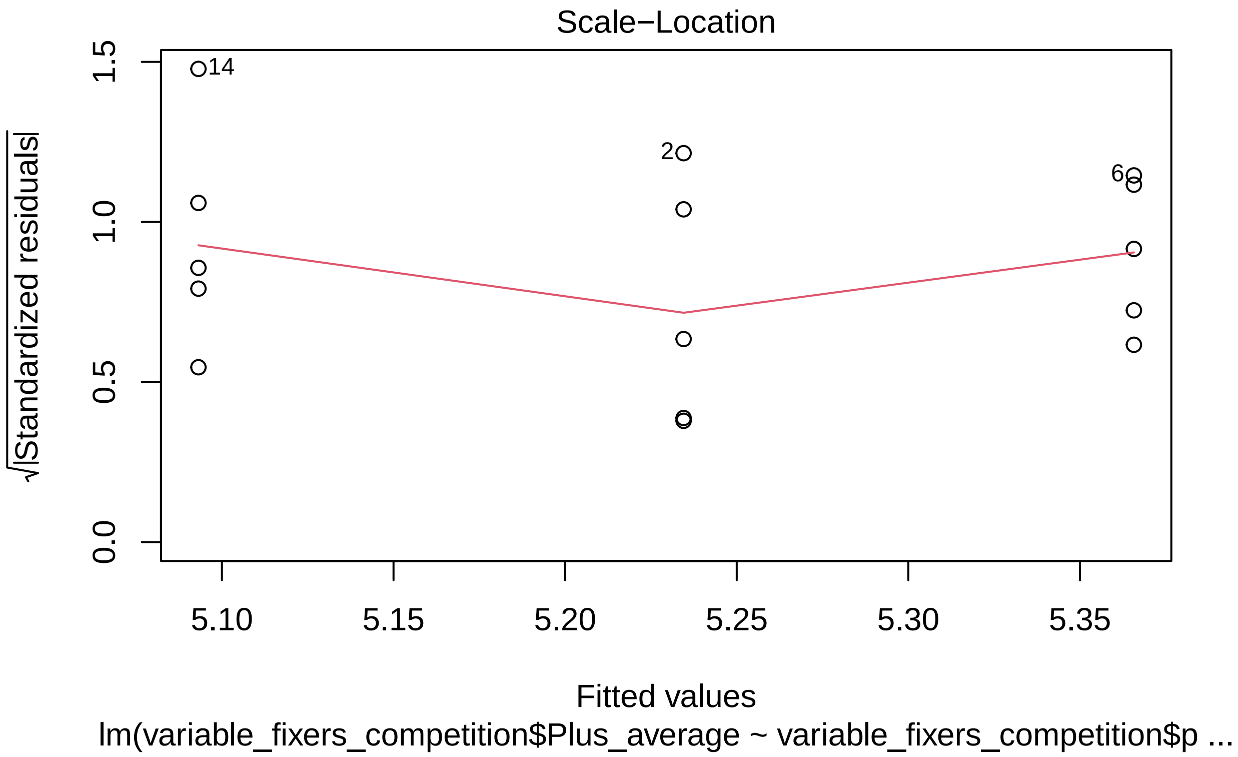
Scale−Location
√|Standardized residuals|
0.0
0.5
1.0
1.5
14
2
6
5.10
5.15
5.20
5.25
5.30
5.35
Fitted values
lm(variable_fixers_competition$Plus_average ~ variable_fixers_competition$p ...

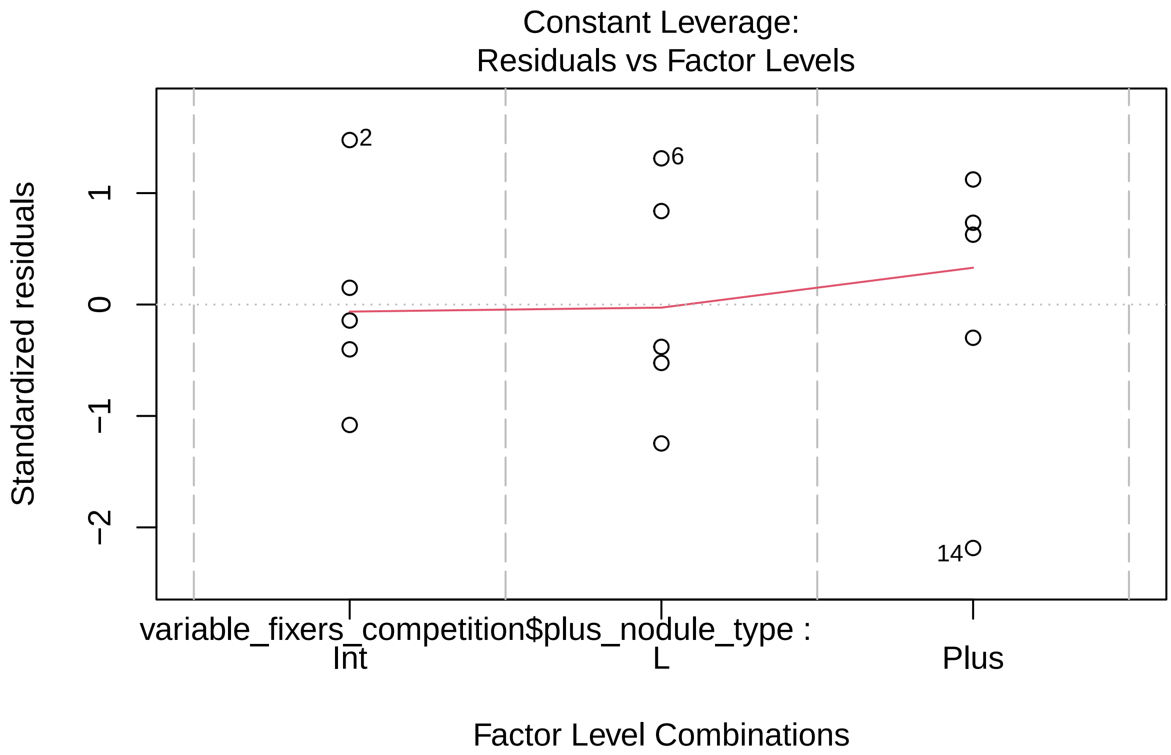


```r
anova(Plus_competition_model)
```

```
## Analysis of Variance Table
##
## Response: variable_fixers_competition$Plus_average
##                                              Df Sum Sq Mean Sq F value Pr(>F)
## variable_fixers_competition$plus_nodule_type  2 0.1858 0.09290  0.1803 0.8373
## Residuals                                    12 6.1843 0.51536
```

```r
plot(L_competition_model)
```

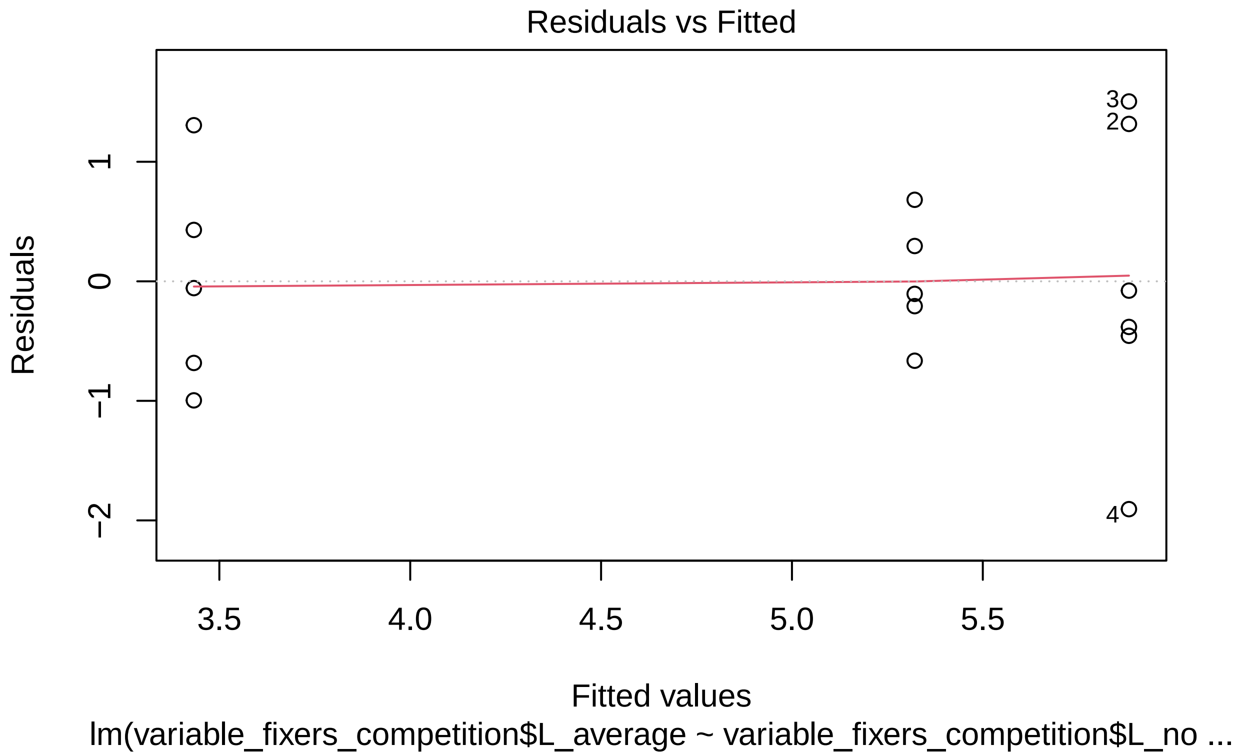
Residuals vs Fitted
Residuals
Fitted values
lm(variable_fixers_competition$L_average ~ variable_fixers_competition$L_no ...
3.5
4.0
4.5
5.0
5.5
1
0
−1
−2
3
2
4

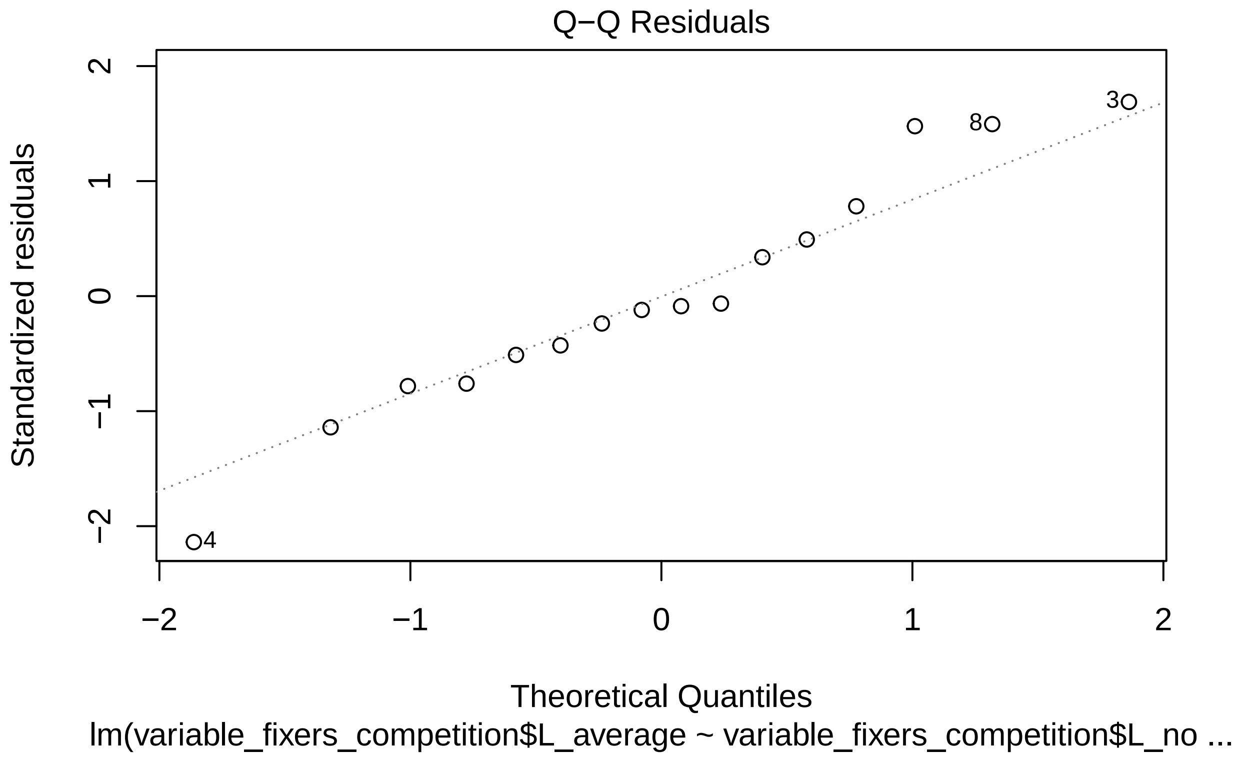

Q−Q Residuals
Standardized residuals
2
1
0
−1
−2
3
8
4
−2
−1
0
1
2
Theoretical Quantiles
lm(variable_fixers_competition$L_average ~ variable_fixers_competition$L_no ...

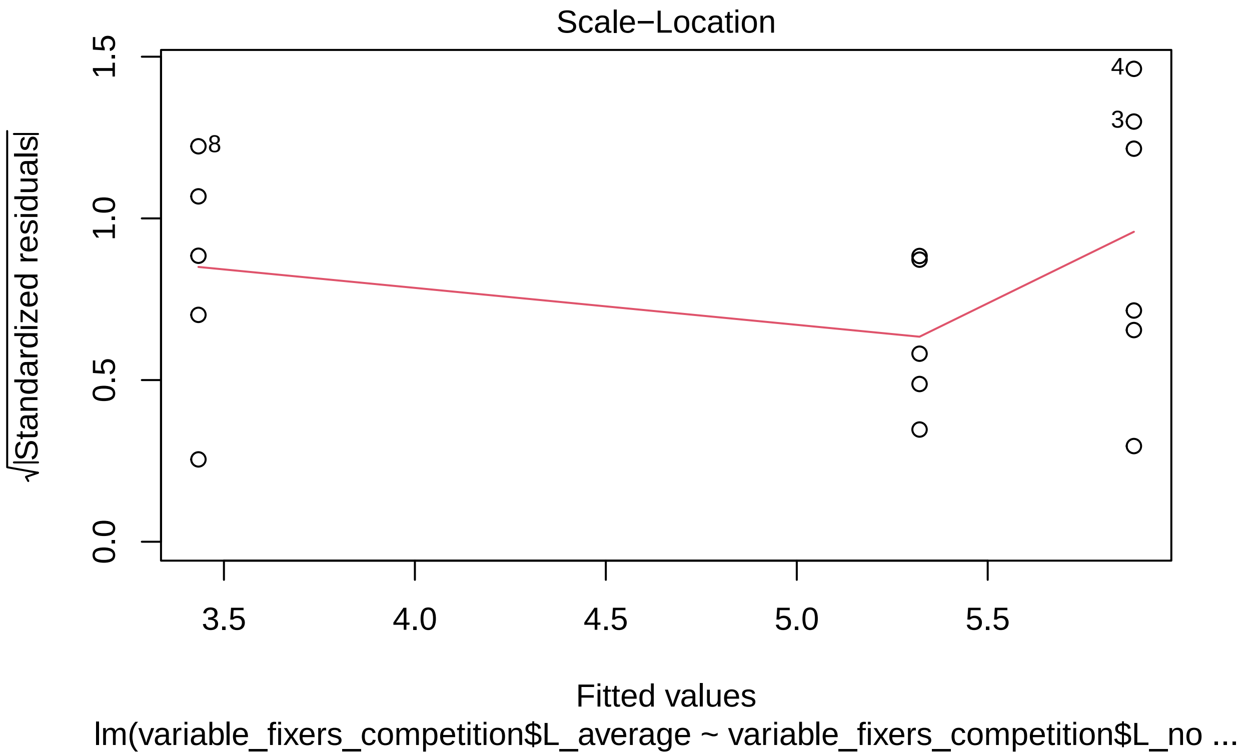
Scale−Location
√|Standardized residuals|
0.0
0.5
1.0
1.5
3.5
4.0
4.5
5.0
5.5
8
4
3
Fitted values
lm(variable_fixers_competition$L_average ~ variable_fixers_competition$L_no ...

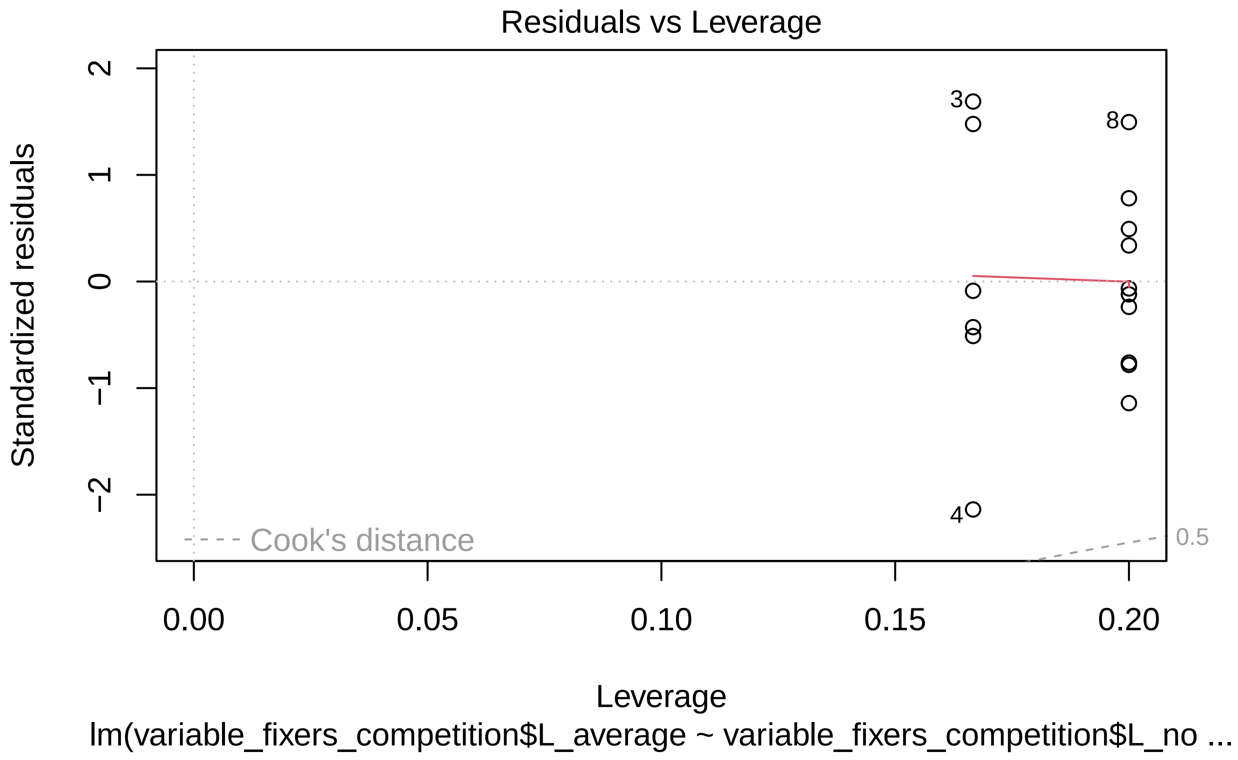


```r
anova(L_competition_model)
```

```
## Analysis of Variance Table
##
## Response: variable_fixers_competition$L_average
##                                           Df Sum Sq Mean Sq F value   Pr(>F)
## variable_fixers_competition$L_nodule_type  2 17.413  8.7066  9.1367 0.003326 **
## Residuals                                 13 12.388  0.9529
## ---
## Signif. codes:  0 '***' 0.001 '**' 0.01 '*' 0.05 '.' 0.1 ' ' 1
```

```r
plot(Int_competition_model)
```

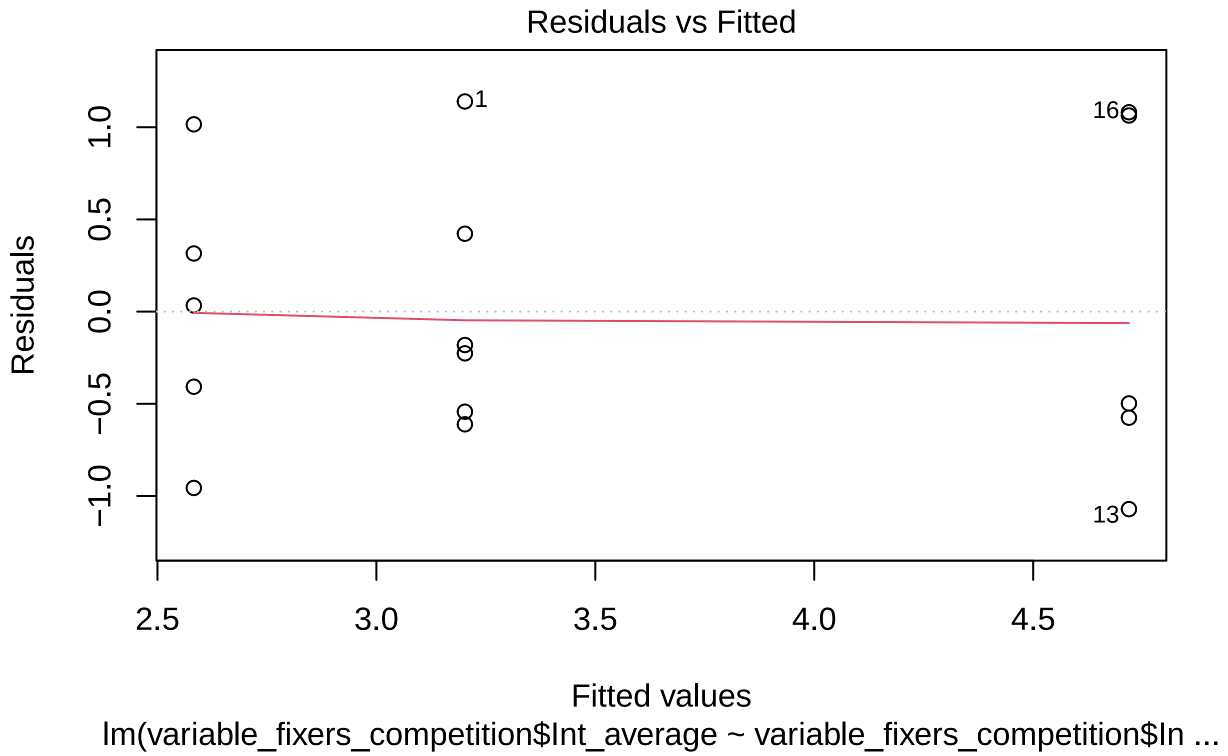
Residuals vs Fitted
Residuals
1.0
0.5
0.0
−0.5
−1.0
1
16
13
2.5
3.0
3.5
4.0
4.5
Fitted values
lm(variable_fixers_competition$Int_average ~ variable_fixers_competition$In ...

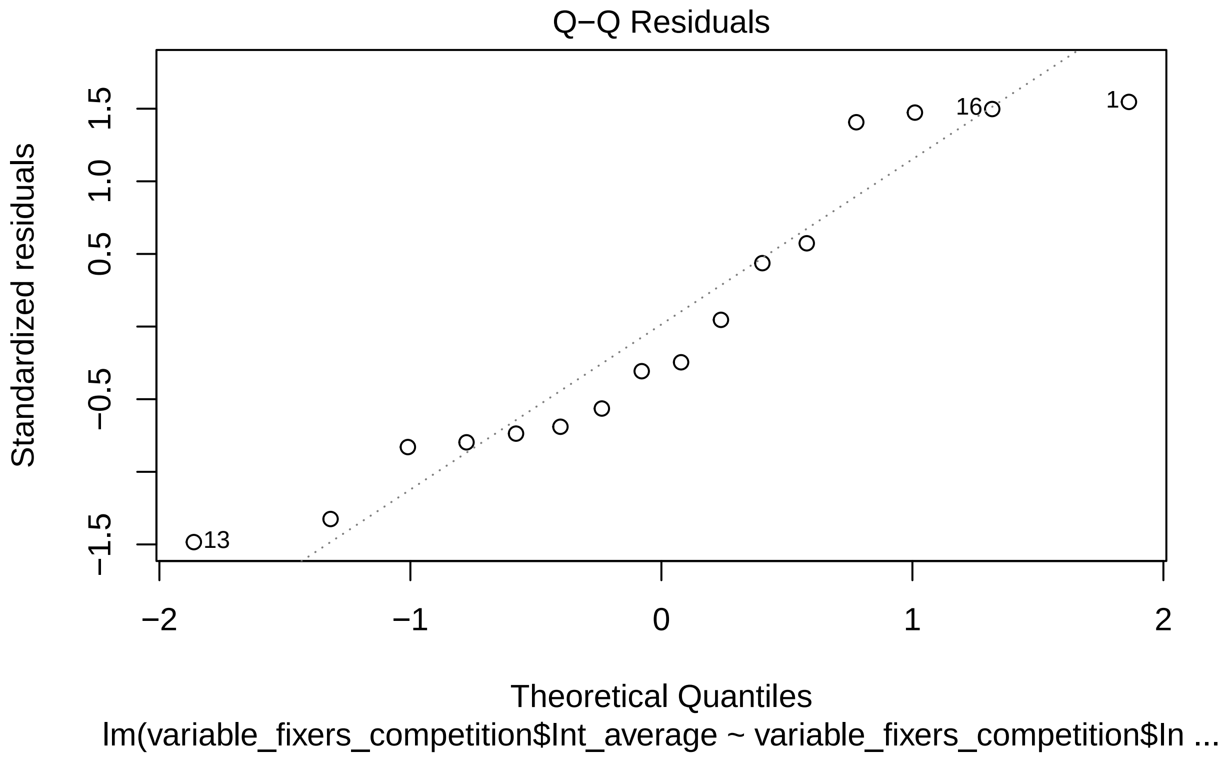
Q−Q Residuals
Standardized residuals
1.5
1.0
0.5
−0.5
−1.5
16
1
13
−2
−1
0
1
2
Theoretical Quantiles
lm(variable_fixers_competition$Int_average ~ variable_fixers_competition$In ...

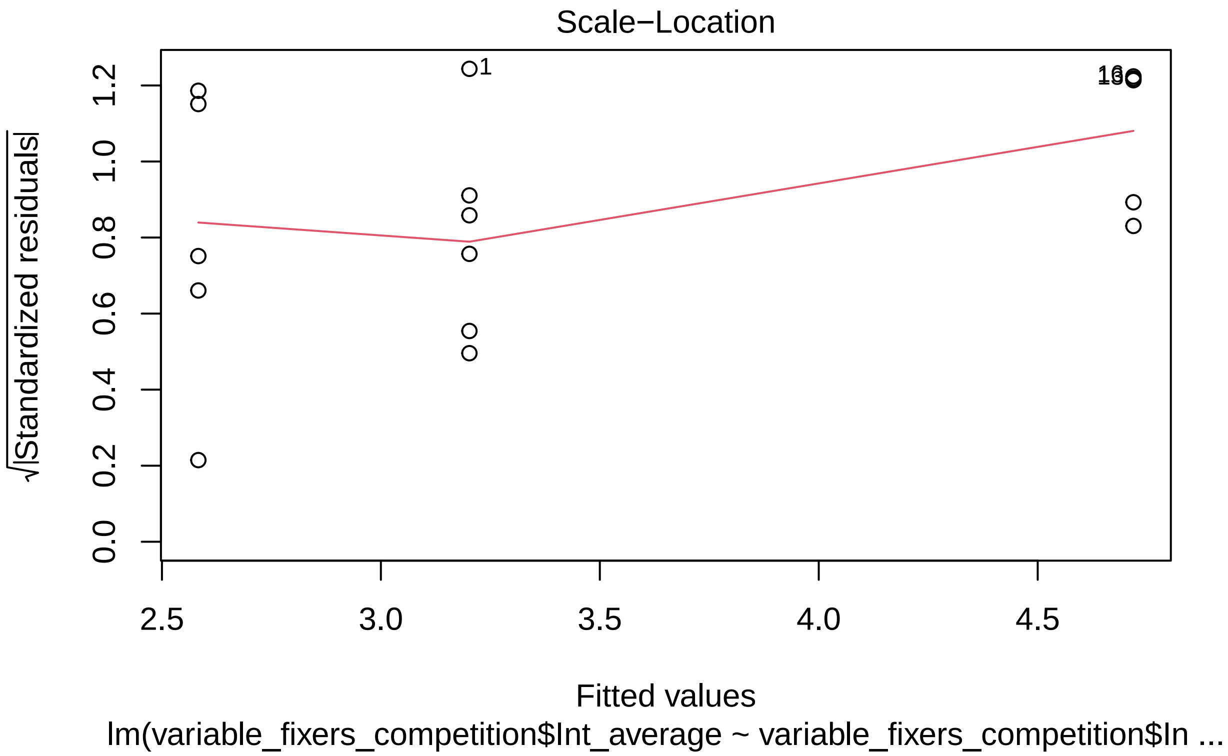
Scale−Location
√|Standardized residuals|
0.0
0.2
0.4
0.6
0.8
1.0
1.2
1
2.5
3.0
3.5
4.0
4.5
Fitted values
lm(variable_fixers_competition$Int_average ~ variable_fixers_competition$In ...

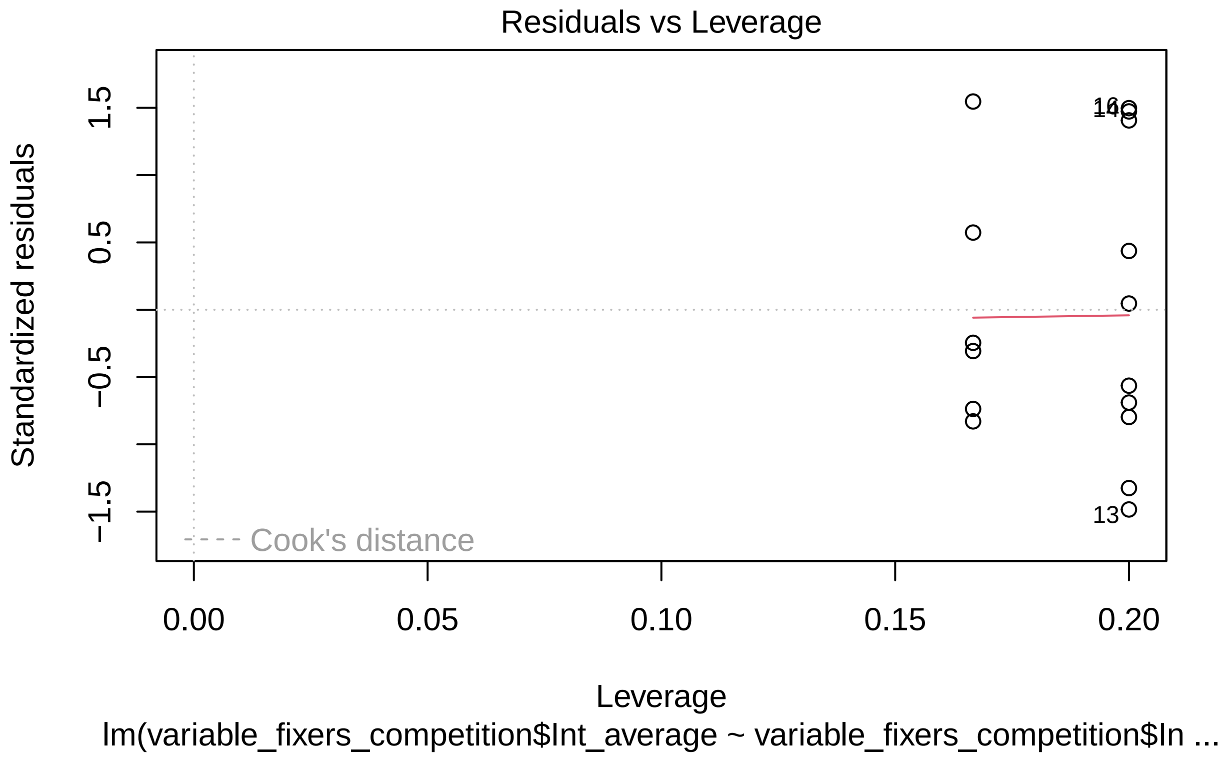


```r
anova(Int_competition_model)
```

```
## Analysis of Variance Table
##
## Response: variable_fixers_competition$Int_average
##                                          Df  Sum Sq Mean Sq F value   Pr(>F)
## variable_fixers_competition$Int_nodule_type  2 12.1566  6.0783  9.3268 0.003075
## Residuals                                13  8.4721  0.6517
##
## variable_fixers_competition$Int_nodule_type **
## Residuals
## ---
## Signif. codes:  0 '***' 0.001 '**' 0.01 '*' 0.05 '.' 0.1 ' ' 1
```

```r
#ANOVA's for int and L nodules showed a
#significant difference but not for the Plus ANOVA
#Therefore Fix Plus nodules did not
#significantly vary in size regardless of co-inoculant

#Therefore we carried out Tukey's post-hoc
#test for our Fix L and Fix int nodules
anova_L <- aov(L_competition_model)

anova_Int <- aov(Int_competition_model)
```

```r
TukeyHSD(anova_L)
```

```
##   Tukey multiple comparisons of means
##     95% family-wise confidence level
##
## Fit: aov(formula = L_competition_model)
##
## $`variable_fixers_competition$L_nodule_type`
##              diff       lwr        upr     p adj
## L-Int    -0.56128 -2.122054  0.9994941 0.6199754
## Plus-Int -2.44960 -4.010374 -0.8888259 0.0030786
## Plus-L   -1.88832 -3.518495 -0.2581446 0.0232071
```

```r
TukeyHSD(anova_Int)
```

```
##   Tukey multiple comparisons of means
##     95% family-wise confidence level
##
## Fit: aov(formula = Int_competition_model)
##
## $`variable_fixers_competition$Int_nodule_type`
##                diff       lwr        upr     p adj
## L-Int    -1.5164883 -2.807221 -0.2257553 0.0213969
## Plus-Int -2.1355600 -3.483687 -0.7874334 0.0028679
## Plus-L   -0.6190717 -1.909805  0.6716613 0.4376085
```

```r
#Fix L nodules were significantly smaller when
#co-inoculated with Fix plus compared to with itself or int

#Fix int nodules were significantly larger when
#co-inoculated with itself compared to with Fix L or PLus
```

We also confirmed the different fixation effectiveness of our strains

```r
#Testing the variation in fixation rates across our strains

Variable_ARA_model <- lm(Variable_ARA$`% of Fix plus`~Variable_ARA$Strain)


plot(Variable_ARA_model)
```

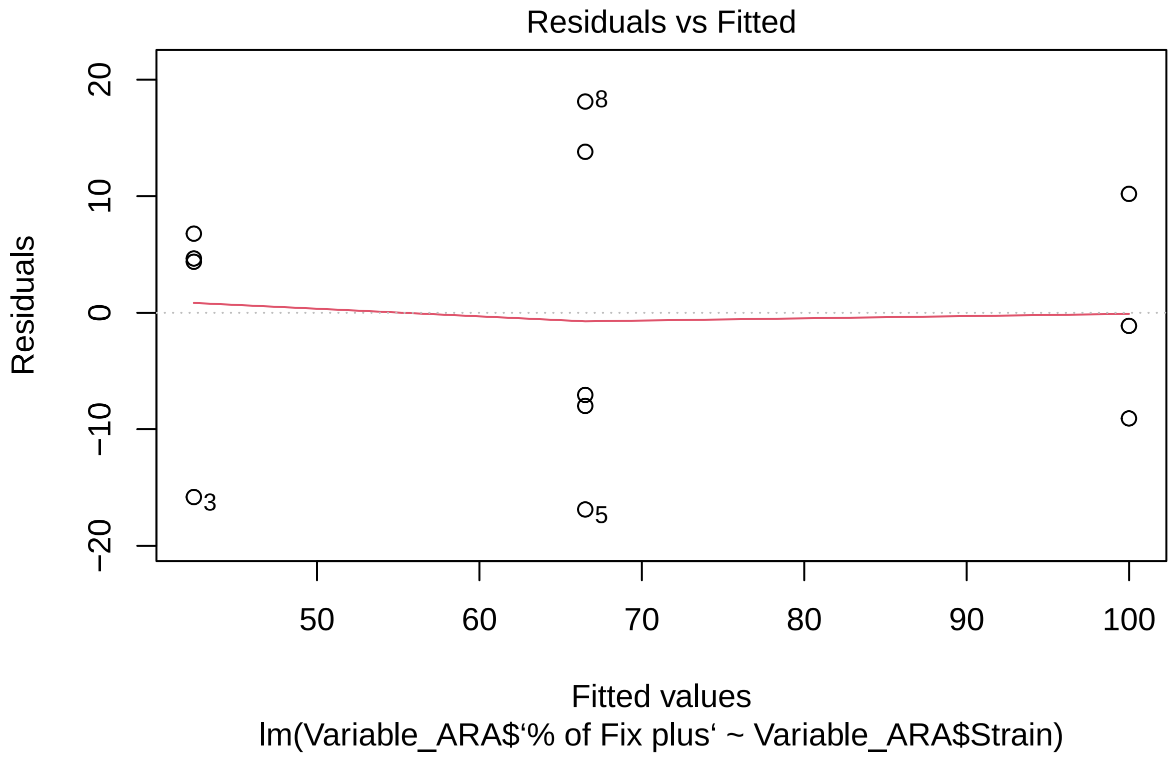
Residuals vs Fitted
Residuals
20
10
0
−10
−20
8
3
5
50
60
70
80
90
100
Fitted values
lm(Variable_ARA$‘% of Fix plus‘ ~ Variable_ARA$Strain)

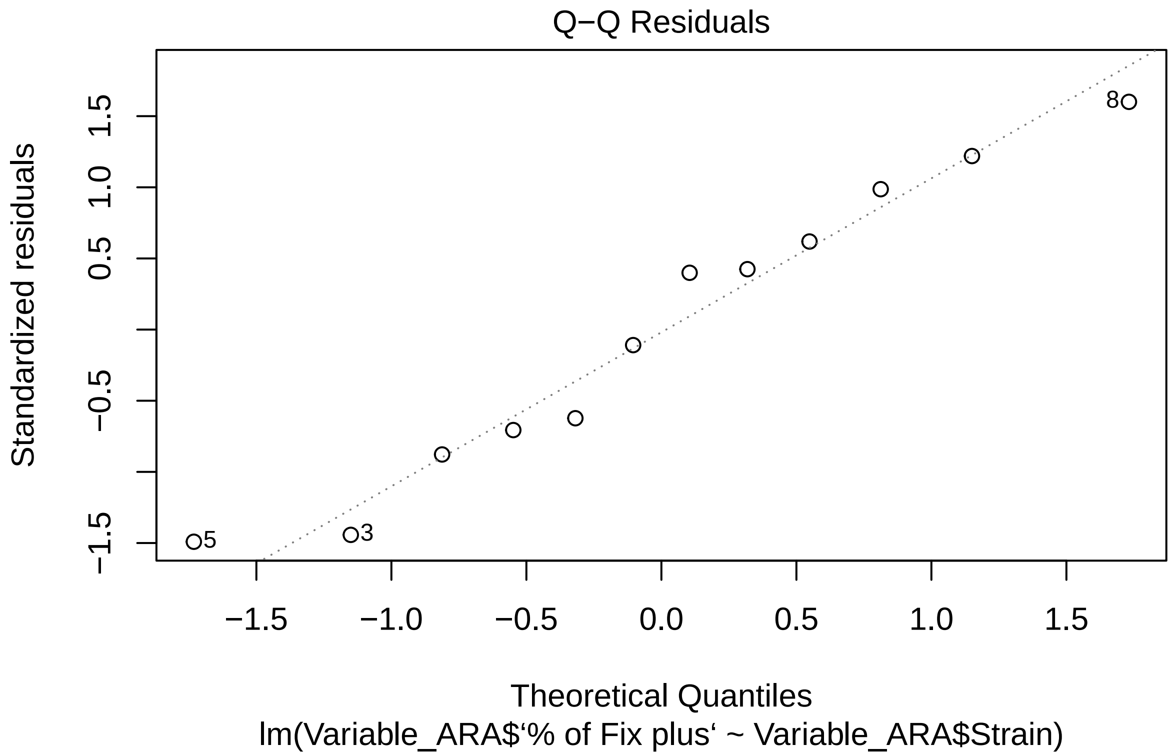
Q−Q Residuals
Standardized residuals
−1.5
−0.5
0.5
1.0
1.5
8
5
3
−1.5
−1.0
−0.5
0.0
0.5
1.0
1.5
Theoretical Quantiles
lm(Variable_ARA$‘% of Fix plus‘ ~ Variable_ARA$Strain)

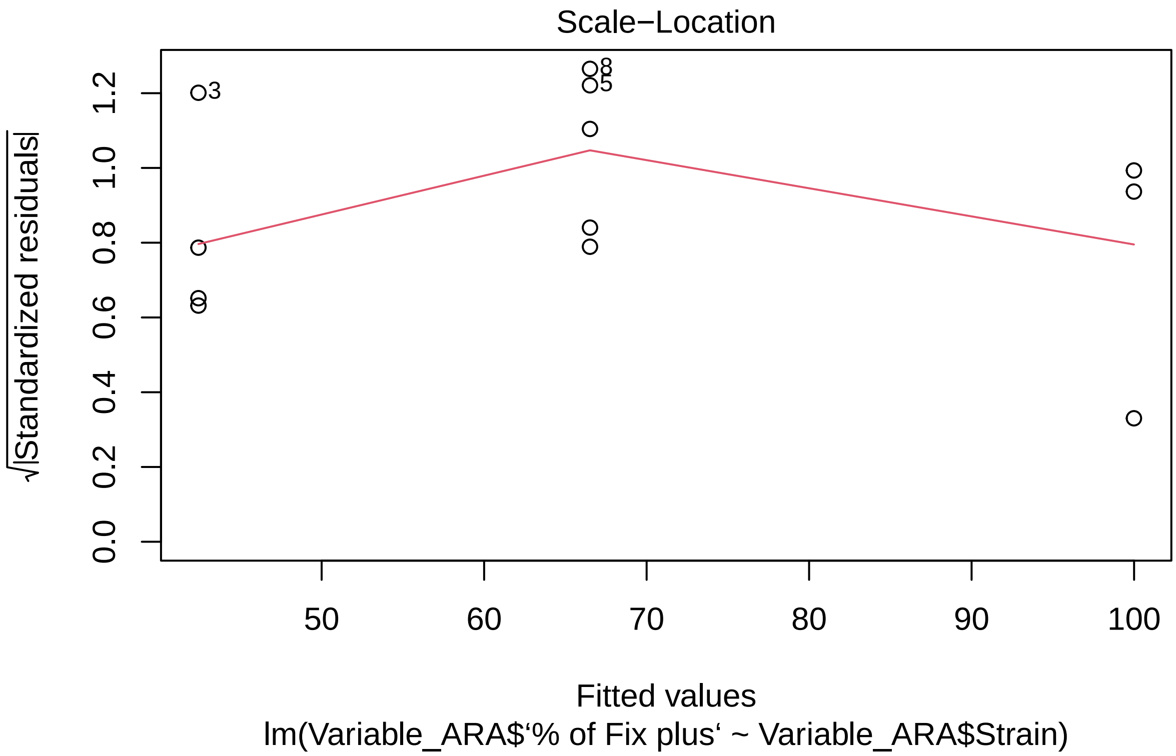
Scale−Location
√|Standardized residuals|
0.0
0.2
0.4
0.6
0.8
1.0
1.2
3
8
5
50
60
70
80
90
100
Fitted values
lm(Variable_ARA$‘% of Fix plus‘ ~ Variable_ARA$Strain)

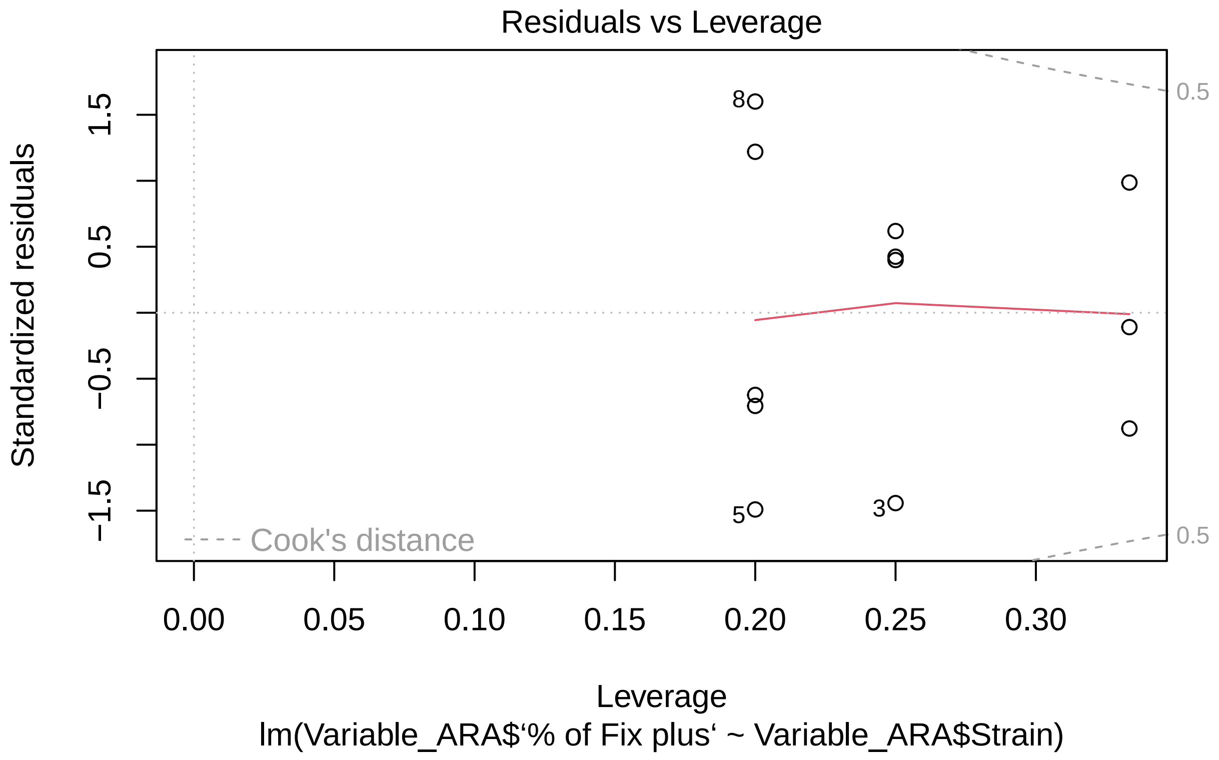


```r
anova(Variable_ARA_model)
```

```
## Analysis of Variance Table
##
## Response: Variable_ARA$‘% of Fix plus‘
##                       Df Sum Sq Mean Sq F value    Pr(>F)
## Variable_ARA$Strain  2 5683.6 2841.78   17.73 0.0007555 ***
## Residuals             9 1442.5  160.28
## ---
## Signif. codes:  0 ’***’ 0.001 ’**’ 0.01 ’*’ 0.05 ’.’ 0.1 ’ ’ 1
```

```r
#ANOVA showed a significant difference so we employed Tukey's post hoc test

Anova_V_ARA <- aov(Variable_ARA_model)

TukeyHSD(Anova_V_ARA)
```

```
##   Tukey multiple comparisons of means
##     95% family-wise confidence level
##
## Fit: aov(formula = Variable_ARA_model)
##
## $‘Variable_ARA$Strain‘
##              diff        lwr      upr     p adj
## L-Int    24.09760  0.3860115 47.80918 0.0465777
```

```
## plus-Int 57.57462 30.5777973 84.57145 0.0005612
## plus-L   33.47703  7.6631243 59.29093 0.0138773
```

```
#Strains were all signficantly different to one another
#in the order Fix Plus > Fix L > Fix Int
```
